# Prognostic significance of *SOCS1* and *SOCS3* tumor suppressors in hepatocellular carcinoma and its correlation to key oncogenic signaling pathways

**DOI:** 10.1101/2020.02.21.958801

**Authors:** Md Gulam Musawwir Khan, Amit Ghosh, Bhavesh Varia, Madanraj Appiya Santharam, Awais Ullah Ihsan, Sheela Ramanathan, Subburaj Ilangumaran

**Affiliations:** Department of Immunology and Cell Biology, Faculty of Medicine and Health Sciences, Université de Sherbrooke, Sherbrooke J1H 5N4, Québec, Canada; CRCHUS, Centre Hospitalier de l’Université de Sherbrooke, Québec, Canada

**Keywords:** hepatocellular carcinoma, SOCS1, SOCS3, TCGA, tumor suppressor, oncogenic signalling, prognosis, precision medicine

## Abstract

Suppressor of cytokine signaling (SOCS) proteins SOCS1 and SOCS3 are considered tumor suppressors in liver hepatocellular carcinoma (LIHC). To gain insight into the underlying molecular mechanisms, the expression of *SOCS1/ SOCS3* was evaluated in The Cancer Genome Atlas LIHC dataset along with key oncogenic signaling pathway genes. *SOCS1* expression was not significantly reduced in HCC yet higher expression predicted favorable prognosis, whereas *SOCS3* lacked predictive potential despite lower expression. Only a small proportion of the cell cycle, receptor tyrosine kinase, growth factor and RAS-RAF-MEK-MAPK signaling genes negatively correlated with *SOCS1* or *SOCS3*, of which even fewer showed elevated expression in HCC and predicted survival. However, many PI3K-AKT-MTOR pathway genes showed mutual exclusivity with *SOCS1*/*SOCS3* and displayed independent predictive ability. Among genes that negatively correlated with *SOCS1*/*SOCS3, CDK2, MLST8, AURKA, MAP3K4* and *RPTOR* showed corresponding modulations in the livers of mice lacking *Socs1* or *Socs3* during liver regeneration and in experimental HCC, and in Hepa1-6 murine HCC cells overexpressing SOCS1/SOCS3. However, Cox proportional hazards model identified *CXCL8, DAB2* and *PIK3R1* as highly predictive in combination with *SOCS1* or *SOCS3*. These data suggest that developing prognostic biomarkers and precision treatment strategies based on *SOCS1/SOCS3* expression need careful testing in different patient cohorts.

## 1. Introduction

Liver cancer, which arises mainly from hepatocellular carcinoma (HCC), is the fifth most prevalent and the third most lethal cancer worldwide [1]. Despite significant advances in understanding the molecular mechanisms of HCC pathogenesis, therapeutic options remain limited and even the most promising drugs such as Sorafenib show only modest efficacy in clinics [2]. Therefore, new therapies targeting various oncogenic signaling pathways are being developed and some are at various phases of clinical testing [3]. Concurrently, the availability of mouse genetic models and high-quality transcriptomic data from pathology specimens through The Cancer Genome Atlas (TCGA) consortium are fueling efforts to gain a deeper understanding of the molecular heterogeneity in the pathogenesis of HCC in order to identify new therapeutic targets and tailor precision treatment strategies [4-8].

Systematic analysis of the aberrant DNA methylation pattern in a distinct region of chromosome 16 in HCC specimens revealed epigenetic silencing of the gene coding of suppressor of cytokine signaling 1 (SOCS1) in up to 65% of human primary HCC tumor samples [9,10]. The *SOCS3* gene is also repressed in 33% of HCC specimens [11]. SOCS1 and SOCS3 proteins have been extensively studied in immune cells as key regulators of cytokine and growth factor receptor signaling [12]. As physiological hepatocyte proliferation is regulated by cytokines and growth factors, most of which also promote neoplastic growth [13-15], several groups, including our own, studied liver regeneration in mice lacking *Socs1* or *Socs3* and their susceptibility to HCC induced by the hepatocarcinogen diethyl nitrosamine (DEN) [16-19]. These studies reported an increased rate of liver regeneration and heightened susceptibility to DEN-induced HCC in these mice. These findings, in corroboration with clinical data on epigenetic repression of *SOCS1* and *SOCS3* genes in HCC specimens, clearly established non-overlapping tumor suppressor functions of SOCS1 and SOCS3. Moreover, HCC invariably arises in cirrhotic livers, which provides not only an inflammatory environment for hepatocarcinogenesis but also increases the availability of cytokines and growth factors [14]. SOCS1 has been implicated in regulating hepatic fibrogenic response, presumably through regulating cytokine and growth factor signaling in hepatic stellate cells and liver resident and infiltrating immune cells [16,20,21]. SOCS3 may also play similar roles in controlling liver fibrosis [22]. Therefore, SOCS1 and SOCS3 may regulate hepatocyte proliferation directly as well as indirectly by modulating the liver tissue environment.

SOCS1 and SOCS3 share maximum sequence homology and structural similarity among the SOCS family members, yet significantly differ in their ability to control cytokine and growth factor signaling [23]. Whereas SOCS1 controls hepatocyte growth factor (HGF) signaling via the receptor tyrosine kinase MET [24], SOCS3 is essential to control IL-6 signaling and also regulates epidermal growth factor receptor (EGFR) signaling [19]. SOCS1 also regulates the paradoxical oncogenic functions of the cell cycle inhibitor CDKN1A, which generally functions as a tumor suppressor [17]. These findings on cell lines and preclinical animal models imply diverse roles for SOCS1 and SOCS3 in regulating hepatocyte proliferation and neoplastic growth. However, whether these functions are compromised in primary HCC is not yet clear.

To gain a deeper understanding of the non-overlapping tumor suppressor functions of SOCS1 and SOCS3, and to identify the signaling pathways that are aberrantly activated in the absence of SOCS1 or SOCS3, we carried out a systematic analysis on the TCGA dataset on liver hepatocellular carcinoma (TCGA-LIHC) [6]. We evaluated how *SOCS1* and *SOCS3* gene expression correlates with genes implicated in hepatocarcinogenesis, placing emphasis on genes that regulate hepatocyte proliferation and survival. Our findings show that the expression of *SOCS1* and *SOCS3* negatively correlates with several genes in a similar fashion, but also show distinct regulation of a number of genes in several oncogenic signaling pathways. The latter could explain, at least partly, the inability of SOCS3 to compensate for the loss of SOCS1 and *vice versa* in animal models of HCC. We identify *SOCS1* but not *SOCS3* as an independent prognostic factor, whereas both display improved predictive potential when combined with certain key genes of the oncogenic signaling pathways. Besides, our findings reveal important differences between published works on putative HCC biomarkers and the TCGA data, highlighting the need for further studies on prognostic biomarkers for precision HCC therapy.

## 2. Materials and Methods

### 2.1. TCGA-LIHC dataset

The gene expression analysis was performed on the RNAseq data from the TCGA provisional dataset on LIHC generated by the TCGA Research Network (https://www.cancer.gov/tcga) [6]. The provisional TCGA-LIHC cohort contains 442 specimens, of which RNAseq V2 data are available for 373 samples. Within this dataset, fifty samples contained paired tumor and adjacent normal tissues. The gene expression dataset was downloaded from the cBioportal suite for cancer genomics research (https://www.cbioportal.org) and analyzed using various publicly available tools as illustrated in the workflow in Supplementary Figure S1.

### 2.2. Correlation between SOCS1/SOCS3 and oncogenic signaling pathway genes

The various oncogenic signaling pathway genes found to be commonly affected in diverse cancers have been identified and categorized by the TCGA working groups [29]. Among these pathways, those related to cell survival and proliferation were chosen for comparative analysis with *SOCS1* and *SOCS3* genes. These pathways include cell cycle control (34 genes), RTK signaling and angiogenesis (19 genes), other growth/proliferation signaling and telomerase (11 genes), RAS-RAF-MEK-MAPK signaling (26 genes) and PI3K-AKT-MTOR signaling (17 genes). The genes within each pathway are listed in the respective figures. Correlation between the expression of *SOCS1*/*SOCS3* and those of the query genes in the aforementioned oncogenic signaling pathways was evaluated by Pearson’s nonparametric correlation analysis (one-tailed) using the GraphPad Prism (version 8) software. The correlation coefficient (ρ-value) was represented in a heatmap to reveal the relationship between *SOCS1/SOCS3* and genes within the selected pathways. Statistical significance of the correlation is indicated asterisks within the heatmap.

### 2.3. Impact of gene expression on patient survival

Correlation between gene expression and patient survival was analyzed using TCGA Clinical Data Resource (TCGA-CDR) available through the UALCAN platform (http://ualcan.path.uab.edu/index.html) [104,105]. UALCAN was used to determine the expression of the query genes in tumor vs non-tumor tissues and across the tumor grades, and its relationship to patient survival. The Kaplan-Meier survival plots were generated by comparing the high expression cases (top 25%) with moderate/low expression (the remaining 75%). Significance of the survival impact in these two groups was measured by log-rank (Mantel-Cox) *p*-values, or by Gehan-Breslow-Wilcox test where indicated.

### 2.4. STRING analysis

The STRING database (http://www.stringdb.org/) was used to illustrate the protein-protein interaction network between SOCS1, SOCS3 and the query proteins in each oncogenic signaling pathway. Only medium and high confidence interactions with a score above 0.40, supported by experiments and/or curated databases on signal transduction pathways were considered for data interpretation.

### 2.5. Cox proportional hazard model

The expression levels of all genes in the selected oncogenic signaling pathways were dichotomized according to the pre-determined cut-off values of low or high expression (≤25^th^ percentile and ≥25^th^ percentile) and the remaining (>75^th^ percentile and <75^th^ percentile). Each list was combined with the dichotomized lists for *SOCS1* and *SOCS3*, resulting in four different dichotomous combinations (low *SOCS1* + low *Gene-X* versus rest, low *SOCS1* + high *Gene-X* vs rest, high *SOCS1* + low *Gene-X* vs rest, high *SOCS1* + high *Gene-X* vs rest). All possible combinations of *SOCS1* with all query genes were entered into a Cox proportional hazards model using the SAS software v9.4 (SAS Institute Inc., Cary, NC). A stepwise selection was used to determine the most predictive combination for patient survival (better or poor survival). The significant effects of the selected combination variables were then validated with a univariate log-rank test for the query gene. The same procedure was applied to *SOCS3*.

### 2.6. Mice studies

Hepatocyte-specific SOCS1-deficient mice, generated by crossing *Socs1*^*fl/fl*^ mice with albumin-Cre (*Alb*^*Cre*^)mice, have been already described [17]. *Socs3*^*fl/fl*^ mice were purchased from the Jackson laboratories (B6;129S4-*Socs3*^*tm1Ayos/J*^) and hepatocyte-specific SOCS3-deficient mice were generated by crossing them with *Alb*^*Cre*^ mice. All animal experiments were carried out with the approval of the Université de Sherbrooke Ethical committee on animal experimentation (protocol number 226-17B) in accordance with the guidelines set by the Canadian Council on Animal care (CCAC). Partial hepatectomy was carried out on 8-10 weeks old mice as detailed previously [28], and remnant liver tissues were harvested after 24h. Experimental HCC was induced by the administration of diethyl nitrosamine to 2-weeks old male pups as previously described [17]. The mice were euthanized after 8 months, and macroscopic liver tumor nodules and adjacent normal tissues were resected. Small pieces of tissues were immersed in RNAlater (ThermoFisher) and stored at -20°C for gene expression analysis.

### 2.7. Cell lines

Murine HCC cell line Hepa1-6 (ATCC: CRL-1830) stably expressing SOCS1 has been previously reported [28]. Hepa1-6 cells were transfected with SOCS3 expression construct in pcDNA3 vector and stable SOCS3 expressing stable lines were tested for the attenuation of IL-6-induced STAT3 phosphorylation.

### 2.8. Quantitative RT-PCR

Total RNA was isolated from liver tissues and cell lines using RiboZol^TM^ (AMRESCO, Solon, OH). After verifying the RNA quality by UV absorption, the first complementary strand was made from 1 µg total RNA using QuantiTect® reverse transcription kit (Qiagen). RT-PCR for the for gene expression analysis was carried out using the CFX-96 thermocycler (Bio-Rad, Mississauga, ON) using the following primers: Mouse *Cdk2* (NM_016756): Fw-GCATTCCTCTTCCCCTCATC; and Rv-GGACCCCTCTGCATTGATAAG; *Aurka* (NM_011497): Fw-TGAGTTGGAAAGGGACATGG and Rv-GGGAACAGTGGTCTTAACAGG; Human *Mlst8* (NM_019988): Fw-CTGAGTCTTCCATCACGTCTG and Rv-GATCTTGGTCTTAGGGATGAGC; *Rptor* (NM_028898): Fw-CACTCCTTGTCTTCATCTGGG and Rv-TGTCATGGTCCTATGTTCAGC; House-keeping genes: mouse *m36b4*; m36B4 (NM_007475.5): Fw-TCTGGAGGGTGTCCGCAAC and Rv-CTTGACCTTTTCAGTAAGTGG.

All primers showed more than 90% efficiency with a single melting curve. Expression levels of the housekeeping gene were used to calculate fold induction of the specific genes modulated by the absence or presence of SOCS1 or SOCS3.

## 3. Results

### 3.1. Reduced Expression Of SOCS1 But Not SOCS3 Correlates With Poor Patient Survival

Analysis of the TCGA-LIHC RNAseq data was carried out following the workflow illustrated in Supplementary Figure S1 and detailed in the methods sections. Contrary to the reports on epigenetic *SOCS1* gene repression in HCC [9,11], the TCGA dataset did not show any significant difference in *SOCS1* expression between tumor tissues and adjacent normal tissues, whereas *SOCS3* expression was significantly reduced in tumor tissues (Figure 1A). Extending this analysis to TCGA-LIHC samples grouped according to the tumor grade also showed no significant difference for *SOCS1* expression across the tumor grades, whereas SOCS3 was significantly reduced with increasing tumor grade (Figure 1B). On the other hand, higher *SOCS1* expression correlated positively with overall patient survival, whereas the *SOCS3* expression level did not correlate with disease outcome (Figure 1C). These data suggest that despite the lack of correlation with the tumor stage, reduced *SOCS1* expression in tumor tissues has an independent prognostic value, whereas reduced *SOCS3* expression *per se* does not have a prognostic significance. Nonetheless, compelling evidence for the non-overlapping tumor suppressor functions of SOCS1 and SOCS3 from genetic ablation studies in mouse models [16-19] prompted us to investigate the relationship between the expression levels of *SOCS1* and *SOCS3* in the TCGA-LIHC dataset and the key signaling pathway genes implicated in carcinogenesis.

**Figure 1.**
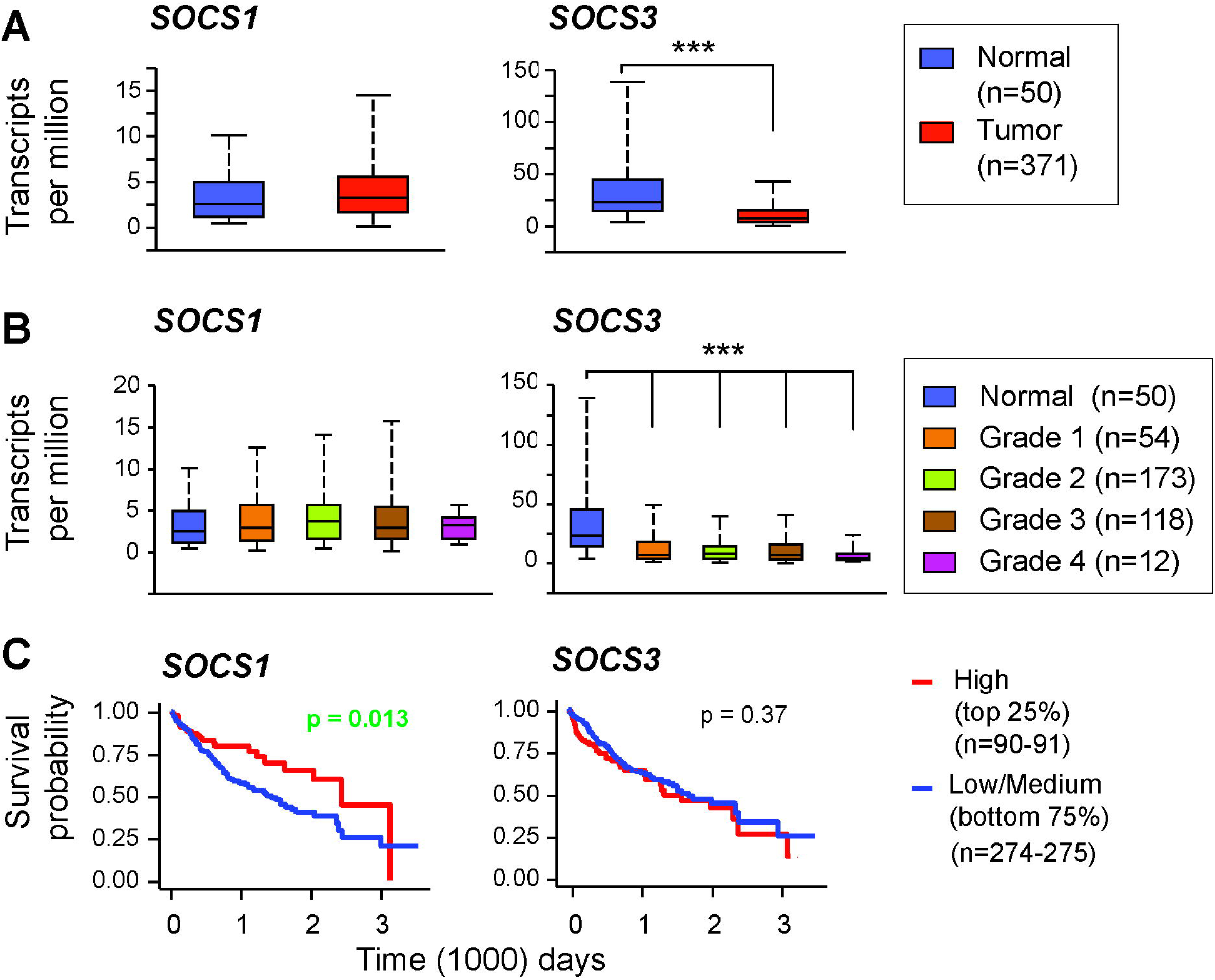
Expression of *SOCS1* and *SOCS3* genes in TCGA-LIHC dataset and their prognostic significance. (A, B) Expression levels of *SOCS1* and *SOCS3* genes in the HCC tumors compared to normal liver tissue in the TCGA dataset. (B) Expression levels of *SOCS1* and *SOCS3* in different grades of HCC specimens. (C) Impact of high *SOCS1* and *SOCS3* expression on overall patient survival. The upper high expression quartile was compared with the remaining three-quarts of low/medium expression in the Kaplan-Meier plot.

### 3.2. Cell Cycle Regulation

SOCS1 and SOCS3 are implicated in the regulation of HGF and IL-6 signaling pathways that promote hepatocyte proliferation and HCC pathogenesis [19,24-28]. Besides, the cell cycle pathway contains the most frequently altered genes in the TCGA-LIHC dataset [29]. Therefore, we evaluated the relationship between the expression of *SOCS1* and *SOCS3* with the thirty-four genes of the cell cycle identified by the cBioportal to be generally deregulated in TCGA datasets of different cancers. STRING analysis predicted these proteins to be tightly interconnected and to *SOCS1* and/or *SOCS3* either directly or indirectly via other members of this group (Supplementary Figure S2A). Next, we retrieved the mRNA expression data for these genes from the TCGA-LIHC dataset and performed non-parametric Spearman correlation (one-way) analysis to determine their common and differential relationship to *SOCS1* and *SOCS3* in terms of co-occurrence or mutual exclusivity.

A heatmap depicting the correlation between the select cell cycle genes with *SOCS1* or *SOCS3* is shown in Figure 2A with the Spearman’s rank correlation coefficient (ρ) aligned at the extremities for *SOCS1*. The ρ-value of -1 and 1 (green to red) implies a stronger linear relationship of mutual exclusivity and co-occurrence, respectively. The significance of the correlation (*p*-value) is depicted within the heatmap. Surprisingly, out of the thirty-four cell cycle genes implicated in oncogenesis, eighteen positively correlated with *SOCS1* in the HCC tumors and fifteen with *SOCS3*, presumably reflecting their induction by growth stimulatory signals such as STAT activation. This notion is supported by a strong positive correlation for both *SOCS1* and *SOCS3* with *STAT3* and *STAT5A* (Figure 2A). It is noteworthy that SOCS1 and SOCS3 were discovered as STAT-inducible proteins [30-32]. High expression of many genes that showed a positive correlation with *SOCS1*, as well as a number of genes that showed no correlation, displayed the ability to independently predict poor prognosis (Supplementary Table 1).

**Figure 2.**
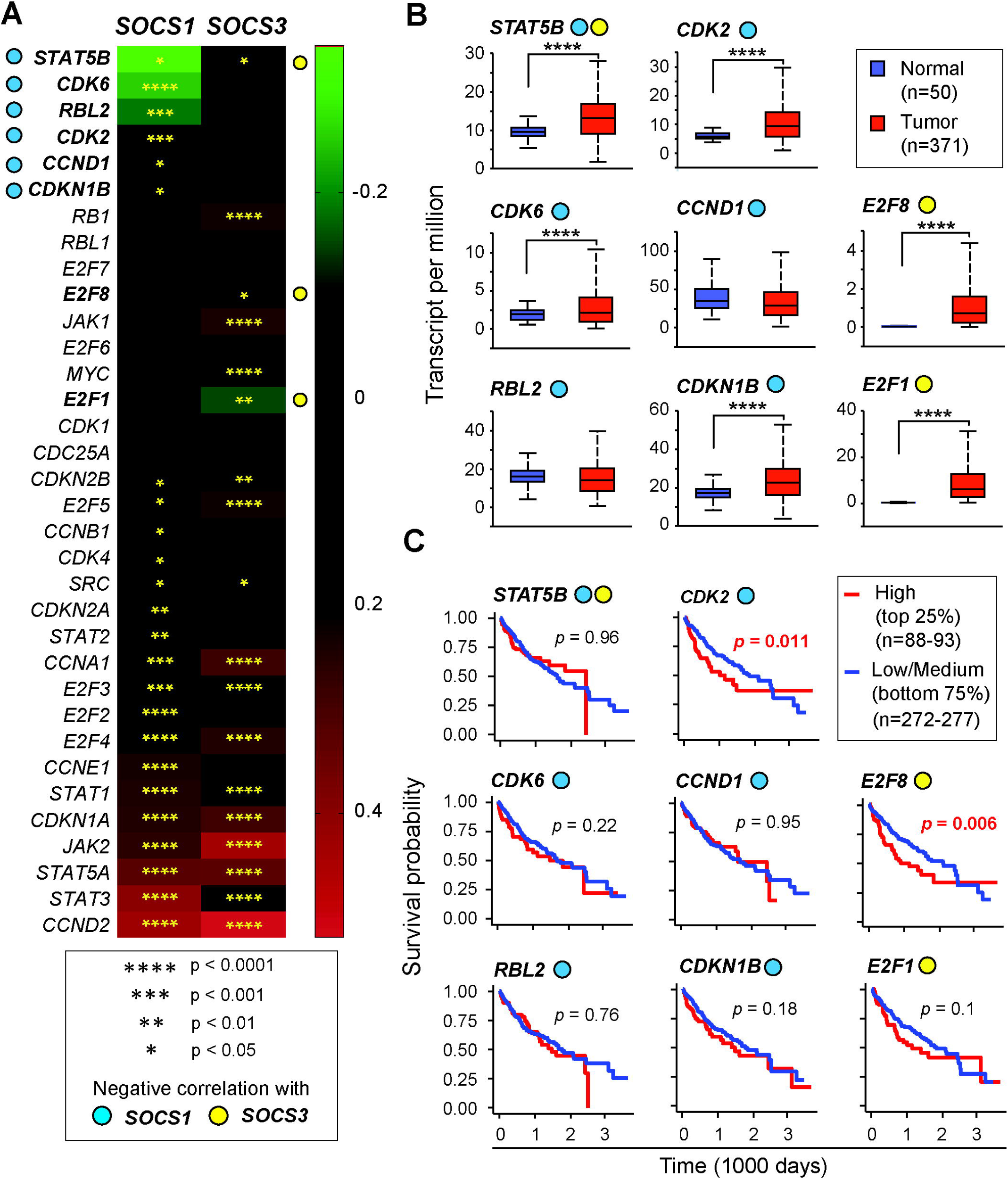
Cell cycle regulation genes in the TCGA-LIHC dataset: correlation with *SOCS1* and *SOCS3* genes, expression in tumors and predictive value. (A) The mRNA expression levels of thirty-four genes of the cell cycle regulation implicated in oncogenesis were evaluated for correlation to *SOCS1* and *SOCS3* expression. The heatmap shows negative (mutual exclusivity) and positive (co-expression) correlations indicated by the color scale on the right. Asterisks within the heatmap indicate the statistical significance of the Spearman correlation. Blue circles on the left indicate Genes showing statistically significant negative correlation with SOCS1 and yellow circles on the right mark genes showing mutual exclusivity with SOCS3. (B) Genes that show significant negative correlation with *SOCS1* and/or *SOCS3* were evaluated for their expression levels in HCC tumors compared to normal liver tissues. (C) Prognostic potential of the above genes was evaluated by comparing the upper quartile of high expression against the remaining three-quarts of low/medium expression by Kaplan-Meier plot. For genes showing statistically significant prognostic potential, with high gene expression correlating poor overall survival, the *p*-values are indicated in red color font.

As the focus of the present study is to understand the impact of the loss of SOCS1 and SOCS3 tumor suppressors on oncogenic signaling, we focused primarily on tumor-promoting genes that showed a negative correlation with *SOCS1* and *SOCS3* expression. Whereas *SOCS1* showed a significant negative correlation with six of the thirty-four cell cycle genes (*STAT5B, CDK6, RBL2, CDK2, CCND1* and *CDKN1B*), *SOCS3* showed mutual exclusivity with only three namely, *STAT5B, E2F8*, and *E2F1* (Figure 2A). Of note, *STAT5B* displayed a weak mutual exclusivity with both *SOCS1* and *SOCS3*, whereas *STAT5A* showed a strong co-occurrence. Next, we examined the expression of the eight genes, which showed mutually exclusivity with *SOCS1, SOCS3* or both, in HCC tumors and assessed their relationship to patient survival. Most of these genes (*STAT5B, CDK6, CDK2, CDKN1B, E2F8*, and *E2F1*) showed high mRNA expression in tumor tissues compared to adjacent non-tumor tissues (Figure 2B) with increased expression levels in poorly differentiated grade 3 and grade 4 tumors (Supplementary Figure S2B). However, the elevated expression of most of these genes in HCC tumor tissues did not predict patient survival except *CDK2* and *E2F8*, a higher expression of which was associated with poor survival (Figure 2C).

### 3.3. RTK signaling and angiogenesis pathways

Receptor tyrosine kinase (RTK) signaling activated by growth factors HGF, EGF and IGF, which provide mitogenic signals needed for physiologic hepatocyte proliferation, can become oncogenic in transformed hepatocytes through receptor amplification, increased ligand availability, deregulated control mechanisms and RTK synergy [15,33,34]. In addition, growth factors that signal via RTKs such as FGF, PDGF and VEGF provide mitogenic signals to hepatic stellate cells, fibroblasts and endothelial cells that promote tumor growth and angiogenesis. SOCS1 and SOCS3 regulate HGF receptor MET and EGFR in hepatocytes, and also impact on the hepatic fibrogenic response that can increase ligand availability [19,24,26,28,35]. Therefore, we examined the relationship between *SOCS1/SOCS3* expression and sixteen driver genes of RTKs signaling and six genes of the angiogenesis pathway, three of which overlap with the RTK pathway. Intriguingly, only a few of these genes, *ERBB3, VEGFA* and *CXCL8* displayed the ability to predict the disease outcome (Supplementary Table 1). STRING analysis revealed that *SOCS1* and *SOCS3* are connected to these groups of genes through *KIT, IGF1R* and *EGFR* (Supplementary Figure S3A). *SOCS1* is coordinately regulated with nine of the sixteen genes in the RTKs signaling and four of the six angiogenesis genes (Figure 3A). *SOCS3* showed additional positive correlations in both pathways. A negative correlation was found only for *ERBB2* (also known as EGFR2, HER2) and *KDR* (also known as VGEFR2/FLK1) with both *SOCS1* and *SOCS3*, and additionally for *EGFR* with SOCS1 (Figure 3A). Among these, elevated expression in cancer tissues was observed for *ERBB2* (Figure 3B), but it did not correlate with patient survival (Figure 3C) possibly due to reduced expression in advanced grade tumors (Supplementary Figure S3B).

**Figure 3.**
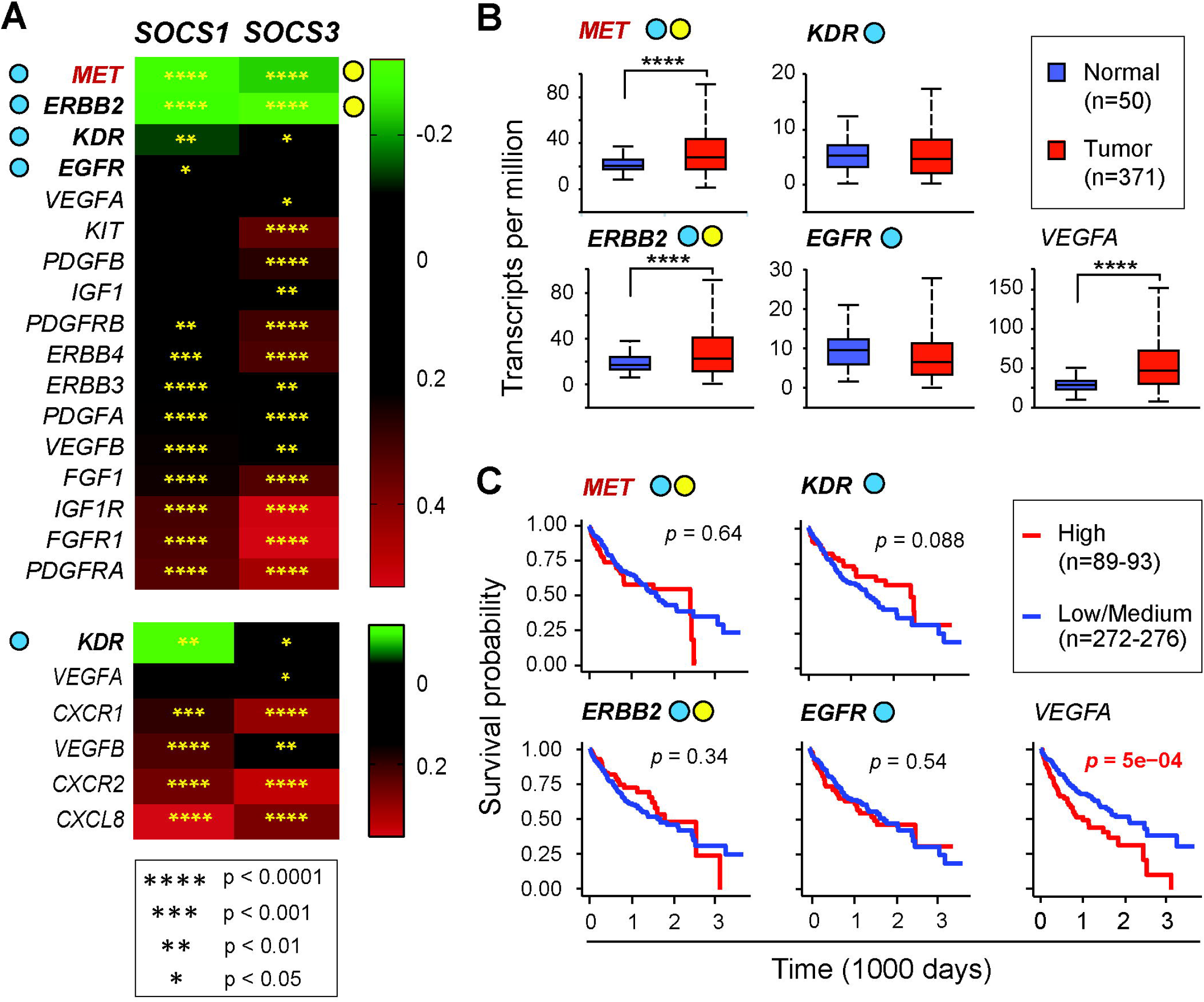
Expression of RTK signaling and angiogenesis pathway genes in the TCGA-LIHC dataset: relationship to *SOCS1* and *SOCS3* expression and survival probability. (A) The mRNA expression levels of sixteen genes of the RTK signaling pathway and six genes of the angiogenesis pathway (two overlapping with the RTK pathway) implicated in oncogenesis were evaluated for correlation to *SOCS1* and *SOCS3* expression as in Figure 3A. MET is not listed in the TCGA oncogenic signaling genes but is included in analysis for reasons detailed in the text. (B) Expression levels of genes, which show significant negative correlation with *SOCS1* and/or *SOCS3*, in HCC tumors and normal liver tissues. (C) Prognostic potential of the above genes.

The HGF receptor MET is an RTK strongly implicated in the pathogenesis of HCC as well as many other cancers, which upregulate MET during disease progression [36]. Others and we have shown that SOCS1 and SOCS3 can regulate MET signaling in the liver and in HCC cell lines [24,26,28]. Deregulation of *MET* through genomic alterations in the TCGA-LIHC dataset has also been reported [6,29]. Therefore, we included *MET* expression in our analysis and found it inversely related to both *SOCS1* and *SOCS3* expression (Figure 3A). Elevated *MET* expression occurs in HCC cancer tissues, but it does not correlate with disease progression or predict patient survival (Figure 3B, 3C; supplementary Figure S3B).

Angiogenesis driven by VEGF is crucial for the progression of HCC, and KDR (Kinase insert Domain Receptor, also known as VEGFR2) is a key signal transducer of VEGFA-induced endothelial cell survival, proliferation, migration and vessel formation [37,38]. The expression of KDR is negatively correlated with SOCS1, but SOCS3 showed a positive correlation with KDR as well as VEGFA (Figure 3A). Whereas VEGFA is elevated in HCC and impacts negatively on patient survival, KDR expression was not increased in HCC (Figure 3B, 3C).

### 3.4. Other growth factors/proliferation signaling pathways and telomerase maintenance

Besides the classical growth factor signaling pathways discussed above, certain non-canonical growth factors and cell proliferation signals contribute to the pathogenesis of several cancers including HCC. This pathway includes genes coding for colony-stimulating factor-1 (CSF1) and its receptor CSF-1R, fibroblast growth factor (FGF)-FGFR and insulin-like growth factor (IGF)-IGFR systems [39-42], and a select set of less well-studied molecules implicated in carcinogenesis such as aurora kinase (*AURKA*) and diphthamide biosynthesis 1 (DPH1) [43,44]. STRING network showed that SOCS1 and SOCS3 associated directly with IGF1R, and SOCS1 additionally with CSF1R (Supplementary Figure S4A). A majority of these eleven genes of this group are coordinately regulated with *SOCS1* and *SOCS3* in the TCGA HCC dataset (Figure 4A, Supplementary Table 1). Two key genes involved in telomerase maintenance reverse transcriptase (*TERT*) and the telomerase RNA component (*TERC*), which are critical for telomerase reactivation during HCC pathogenesis [45], are also included within this group. Notably, significant mutual exclusivity was observed for *SOCS3* with *TERT, TERC* and *AURKA*, and for SOCS1 with *DPH1* (Figure 4A). All these four genes showed higher mRNA expression in HCC tumors compared to normal liver tissue, with high expression of *AURKA* and *DPH1* across the tumor grades (Figure 4B, Supplementary Figure S4B). However, among them, only *AURKA* displayed significant predictive potential with high expression correlating with poor survival (Figure 4C). Even though long telomeres characterize HCC, *TERT* and *TERC* expression levels lacked predictive potential in the TCGA-LIHC dataset.

**Figure 4.**
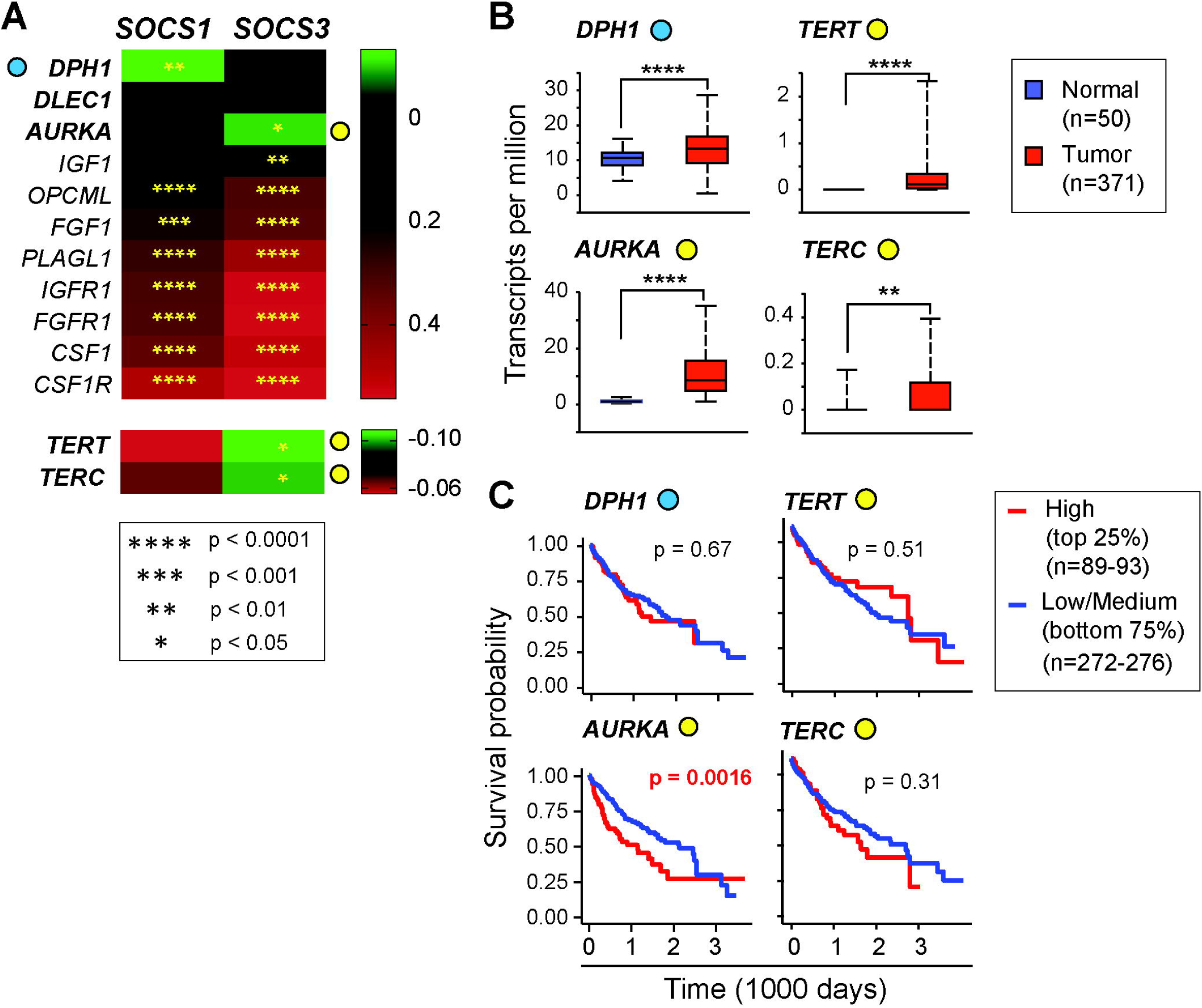
Other growth signaling and telomere maintenance genes: correlation with genes, expression in tumors and prognostic significance in the TCGA data on HCC. (A) Correlation between *SOCS1* and *SOCS3* gene expression with other growth signaling pathway genes implicated in oncogenesis. (B) Expression levels of genes, which show a significant negative correlation with *SOCS1* and/or *SOCS3*, in HCC tumors and normal liver tissues. (C) Predictive value of the above genes.

### 3.5. RAS-RAF-MAPK signaling pathways

The mitogen-activated protein kinases (MAPK) contribute to carcinogenesis via promoting many cellular functions such as cell survival, proliferation and epithelial to mesenchymal transition [46]. This pathway includes extracellular signal-regulated kinase (ERK), c-Jun N-terminal kinase (JNK) and p38 stress-activated kinase (SAPK), of which ERKs activated downstream of growth factor RTKs, and JNKs activated by inflammatory stimuli are strongly implicated in HCC pathogenesis. The canonical MAPK pathway involves activation of the RAS GTPase and RAF kinases, and then sequential activation of MAPKKK (MAP3K) and MAPKK (MAP2K) kinases leading to MAPK activation. Activating mutations of *RAS* and *RAF*, and inactivation/repression of endogenous regulators of RAS such as *RASSF1* and *DAB2* are common in many cancers including HCC [47,48]. STRING analysis showed that SOCS1 is directly connected to this pathway through HRAS and SOCS3 with both HRAS and KRAS (Supplementary Figure S5A). Of the twenty-six oncogenic drivers of this pathway, *SOCS1* showed strong mutual exclusivity with eight genes including *RAF1, BRAF, MAP2K5, MAP3K2, MAPK1* (ERK2), *MAPK6* (ERK3), *MAPK8* (JNK1) and *MAPK14* (p38 SAPK), some of which also showed negative correlation with *SOCS3* (Figure 5A). Many of these negatively correlated genes are highly expressed in HCC across the disease spectrum (Figure 5B, Supplementary Figure S5B), but the expression of only *MAPK1* and *BRAF*, and *MAP3K4* demonstrate the ability to predict patient survival (Figure 5C). Intriguingly, *SOCS1* showed a strong positive correlation with *HRAS* and *SOCS3* with *KRAS*, and both with *RASSF1, DAB2* and *MAPK3* (ERK1) (Figure 5A).

**Figure 5.**
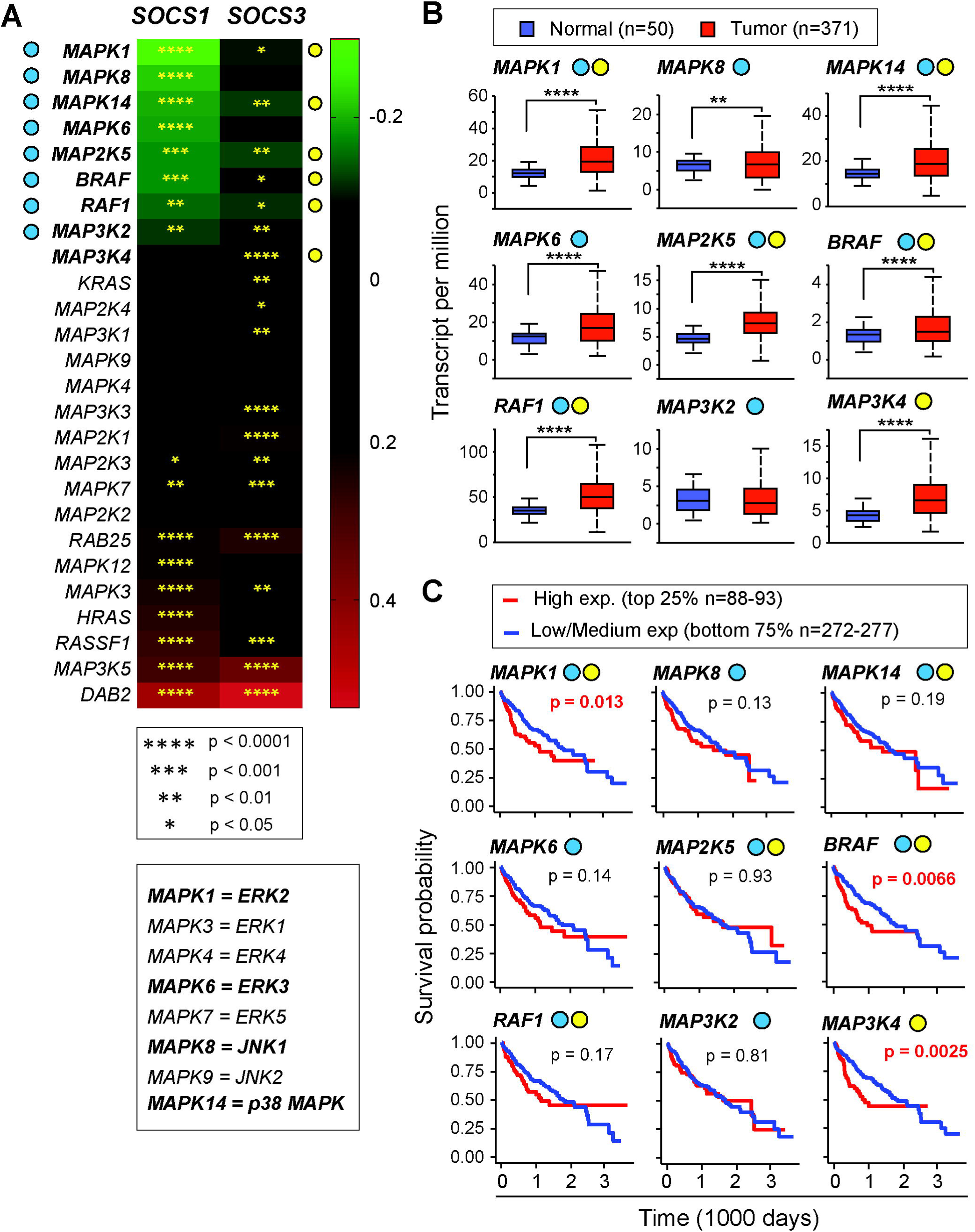
Correlation between *SOCS1, SOCS3* and the RAS-RAF-MEK-MAPK pathway genes and their prognostic significance in the TCGA-LIHC dataset. (A) The oncogenic RAS-RAF-MEK-MAPK pathway genes were compared with SOCS1 and SOCS3 to assess mutual exclusivity and co-expression. Certain common names of genes in this pathway are indicated below. (B) Expression levels of genes, which show significant negative correlation with *SOCS1* and/or *SOCS3*, in HCC tumors and normal liver tissues. (C) Prognostic potential of the above genes.

### 3.6. PI3K-AKT-MTOR Signaling pathway

A key growth-promoting pathway in normal and cancer cells is the PI3K-AKT-MTOR pathway that is deregulated in multiple cancers including HCC and is considered an important target for therapy [49-51]. Activation of the mechanistic target of rapamycin (MTOR, previously called mammalian TOR) occurs via PI3K-AKT pathway downstream of growth factor and cytokine signaling. Therefore, we examined in the TCGA-LIHC dataset the relationship between *SOCS1/SOCS3* expression and a select set of 17 genes of this pathway that are most frequently deregulated in cancers (Figure 6A). STRING analysis revealed that only *SOCS1* is related to this pathway through PIK3R1, the regulatory subunit of PI3K (Supplementary Figure S6A). Nonetheless, 5 of these 17 genes showed a negative correlation with both *SOCS1* and *SOCS3* (*PIK3R1, PDPK1, RPTOR, PTEN, AKT2*), with *PIK3R1* showing the strongest mutual exclusivity. Additionally, *TSC1, MTOR*, and *PIK3CA* revealed a negative correlation only with *SOCS1* whereas *AKT1S1, TSC2*, and *MLST8* showed mutual exclusivity only with *SOCS3*, making PI3K-AKT-MTOR pathway the most closely related to *SOCS1/3* (Supplementary Table 1, Figure 7). Notably, *PIK3CA* (the catalytic subunit of PI3K) showed mutual exclusivity with *SOCS1* but co-occurrence with *SOCS3*, whereas *AKT1S1*, which is a target of AKT, showed an inverse relationship (Figure 6A).

**Figure 6.**
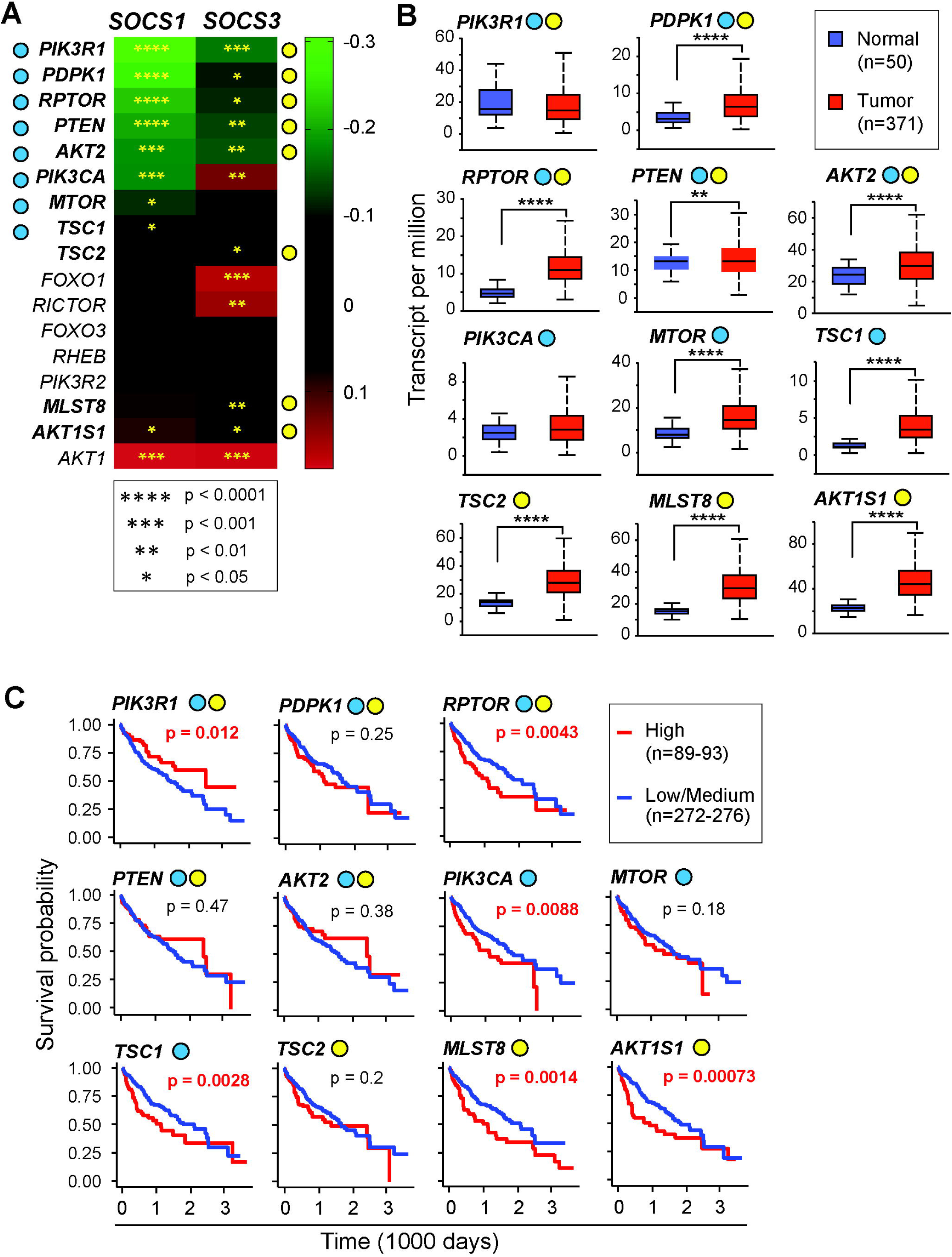
PI3K-AKT-MTOR pathway genes in the TCGA-LIHC dataset: relationship to *SOCS1* and *SOCS3* genes and predictive value. (A) Correlation between *SOCS1* and *SOCS3* gene expression with thePI3K-AKT-MTOR signaling pathway genes implicated in oncogenesis. (B) Expression levels of genes, which show a significant negative correlation with *SOCS1* and/or *SOCS3*, in HCC tumors and normal liver tissues. (C) Predictive value of the above genes. Note that the high expression of *PIK3R1* predicts better prognosis.

**Figure 7.**
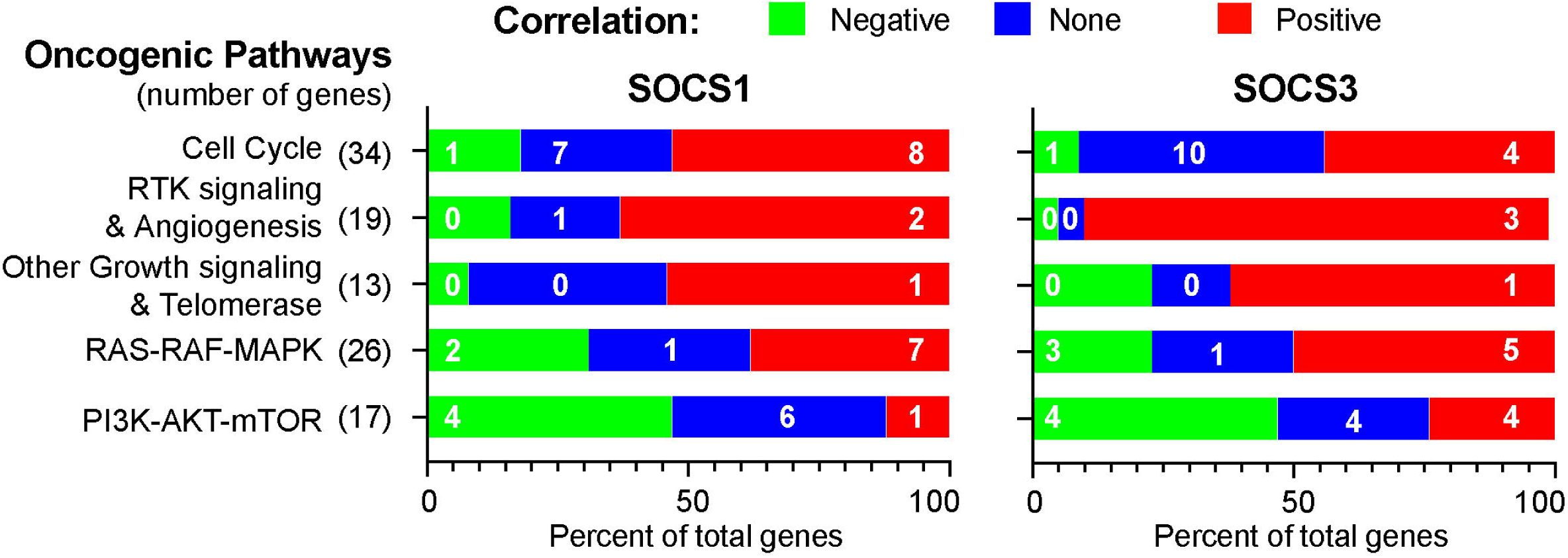
Summary of the correlation between *SOCS1, SOCS3* and the oncogenic signaling pathway genes. Numbers within the bars denote the number of genes with independent prognostic value.

Among the eleven PI3K-AKT-MTOR pathway genes negatively correlated with *SOCS1/SOCS3*, all except *PIK3CA* and *PIK3R1* showed significantly elevated expression in HCC tumors compared to normal liver tissue (Figure 6B, Supplementary Figure S6B). Most of these genes also showed significant predictive value, with high expression associated with poor survival (Figure 6C). Intriguingly, elevated expression of *PIK3R1*, which showed a negative correlation with *SOCS1* and *SOCS3*, was associated with a better disease outcome (Figure 6C).

### 3.7. Predictive potential of oncogenic signaling genes inversely correlated to SOCS1 and SOCS3 and their validation in experimental animal and cellular models

As SOCS1 and SOCS3 are tumor suppressors implicated in regulating cytokine and growth factor signaling pathways, we expected a predominantly inverse correlation between *SOCS1/SOCS3* and oncogenic signaling pathway genes implicated in HCC. However, this was only observed within the PI3K-AKT-MTOR pathway (Figure 7; Supplementary Table 1). Surprisingly, a larger proportion of the oncogenic signaling pathway genes showed a positive correlation with *SOCS1* and/or *SOCS3* (Figure 7; Supplementary Table 1). Among the genes that showed a negative correlation with *SOCS1* and/or *SOCS3*, ten genes displayed independent predictive potential and showed upregulation in tumor tissue, with the exception of *PIK3R1* (Table 1).

**Table 1.**
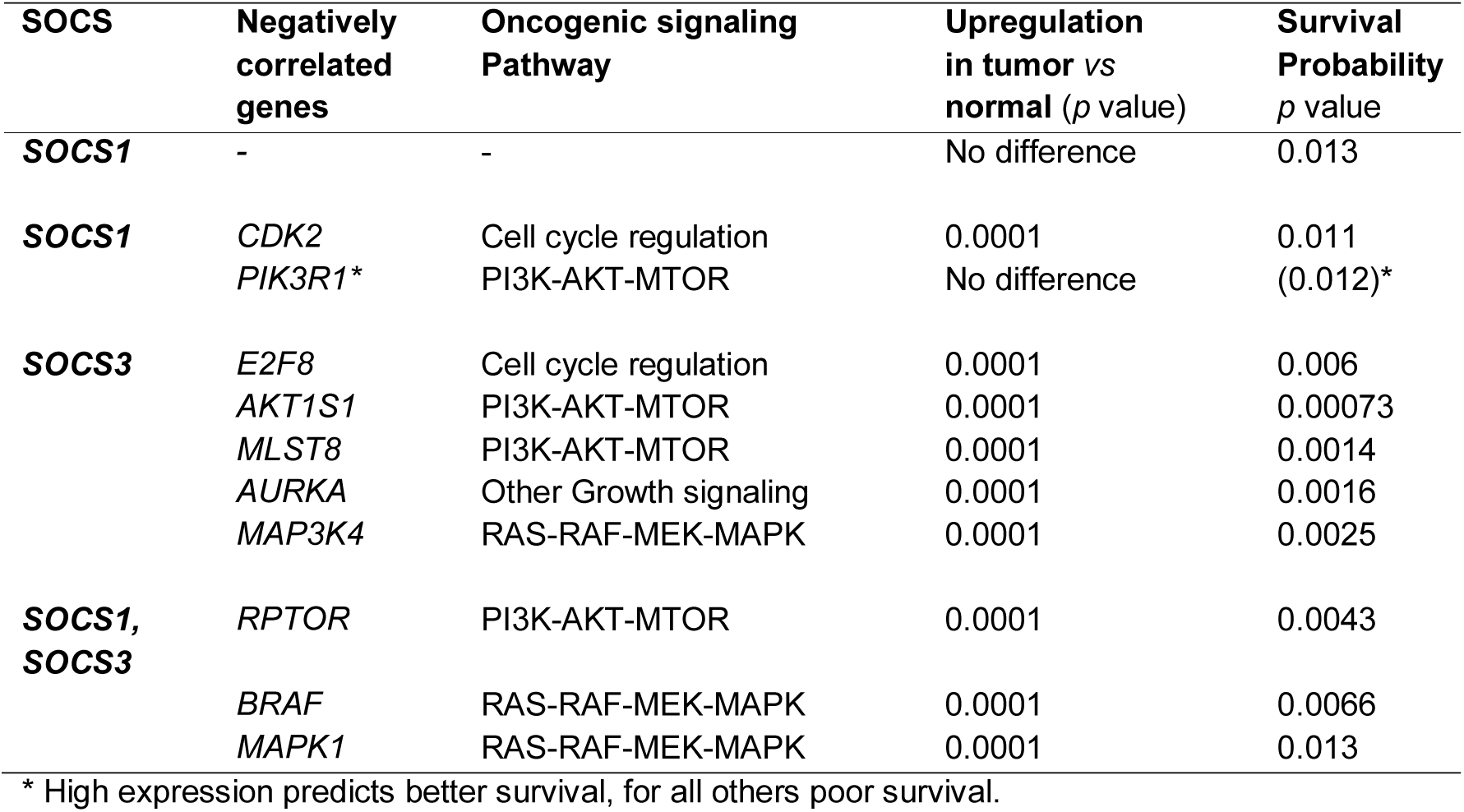
Genes that negatively correlate with *SOCS1* and/or *SOCS3* in the TCGA-LIHC dataset, and their impact on patient survival.

Next, we used mice lacking SOCS1 or SOCS3 in hepatocytes to validate key oncogenic signaling pathway genes that negatively correlated with *SOCS1* or *SOCS3* in the TCGA-LIHC dataset. Unlike the TCGA-LIHC data, wherein the elevated expression of SOCS1 or SOCS3 could result from cells other than hepatocytes, the knockout mice offer the advantage of hepatocyte-specific loss of SOCS1 or SOCS3 expression. Physiological hepatocyte proliferation was induced in *Socs1*^*fl/fl*^*Alb*^*Cre*^, *Socs3*^*fl/fl*^*Alb*^*Cre*^ and *Socs1*^*fl/fl*^*Socs3*^*fl/fl*^ control mice by partial hepatectomy (PH), and gene expression was evaluated in regenerating livers 24h later (Figure 8A). To study pathological hepatocyte proliferation, HCC was induced in hepatocyte-specific SOCS1- or SOCS3-deficient mice using the hepatocarcinogen diethyl nitrosamine (DEN; Figure 8B). Liver tumor nodules and adjacent non-tumor tissues were obtained 8-10 months after DEN injection and were evaluated for gene expression. Additionally, the murine HCC cell line Hepa1-6 stably expressing SOCS1 (Hepa-SOCS1), SOCS3 (Hepa-SOCS3) or control vector (Hepa-vector) were exposed to the inflammatory cytokine IL-6 [25], which is implicated hepatocyte proliferation and hepatocarcinogenesis, and induction of the oncogenic signaling genes was evaluated (Figure 8C).

**Figure 8.**
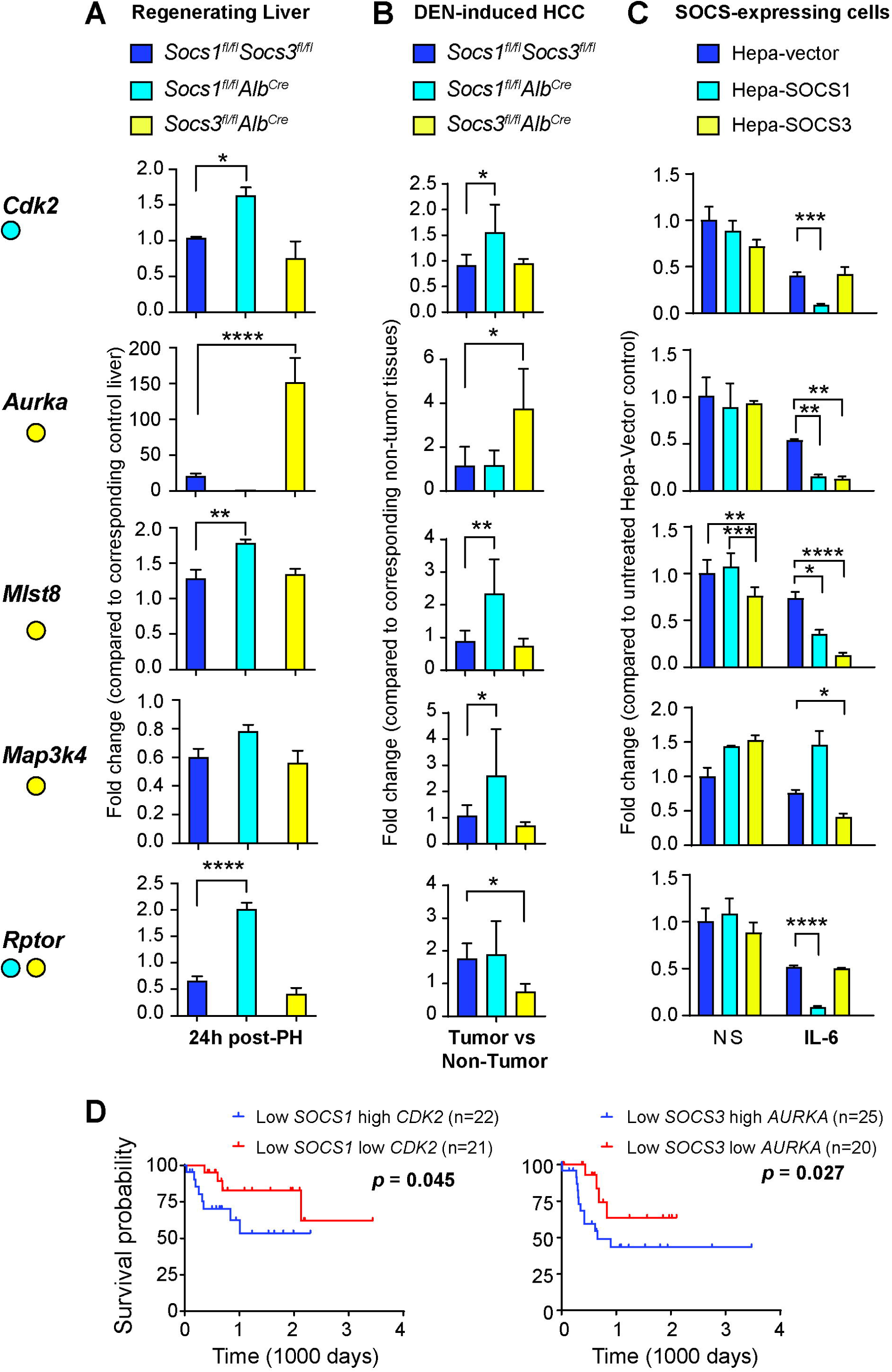
Validation of the oncogenic signaling pathway genes that negatively correlate with *SOCS1* or *SOCS3* in murine and cell line models. (A) Partial hepatectomy was carried out on 8-10 weeks old mice lacking *Socs1* or *Socs3* in hepatocytes and control mice. The expression of the indicated genes in the regenerating livers was evaluated 24h later by qRT-PCR. n=4-6 mice per group. (B) Mice lacking *Socs1* or *Socs3* in hepatocytes and control mice were treated with DEN at 2 weeks of age and livers collected at 8-10 months of age. Tumor nodules and adjacent normal liver tissues were resected and expression of the indicated genes was evaluated by qRT-PCR. n=4-6 mice per group. (C) Murine HCC cell line Hepa1-6 was transfected with SOCS1 or SOCS3 expression constructs or the control vector, and expression of the indicated genes was evaluated 3h after IL-6 stimulation. Data shown are the mean + standard deviation from three independent experiments. *p*-values were calculated by 2-way ANOVA along with Tukey’s Multiple Comparison test: * *p* <0.05, ** *p* <0.01, *** *p* <0.001, **** *p* <0.0001. (D) Prognostic potential of segregating low SOCS1 or SOCS3 expression along with high CDK2 or AURKA expression in the TCGA-LIHC dataset. Kaplan-Meier plot, the number of specimens within each group and the *p*-values calculated by the Gehan-Breslow-Wilcoxon test are shown.

*Cdk2*, which is negatively correlated with *SOCS1* expression in the TCGA dataset, was significantly upregulated in SOCS1-deficient liver following PH as well as in DEN-induced HCC and is downregulated in IL-6-stimulated Hepa-SOCS1 cells. Similarly, *Aurka*, which is negatively correlated with *SOCS3* in the TCGA dataset, was upregulated more than 100-fold in SOCS3-deficient, but not in SOCS1-deficient liver, following PH and in DEN-induced HCC, and is downregulated in IL-6-stimulated Hepa-SOCS3 cells (Figure 8A-C). Loss of SOCS3 in the liver did not affect *Cdk2*, and *Aurka* was not affected by the loss of SOCS1. Strikingly both SOCS3 and SOCS1 diminished the expression of *Aurka* in Hepa cells (Figure 8C). The diminution of *Cdk2* and *Aurka* in control Hepa-vector in cells could be attributed to the induction of endogenous SOCS1, as reported earlier [28]. In contrast to *Aurka, Mlst8* and *Map3k4*, which are negatively correlated with *SOCS3* in the TCGA dataset, showed downregulation in Hepa-SOCS3 cells but were not affected by SOCS3 deficiency in the regenerating livers or in HCC tissues, although discernible upregulation was observed in the absence of SOCS1 (Figure 8A-C). *RPTOR*, which is negatively correlated with both *SOCS1* and *SOCS3* in the TCGA dataset, was increased in the regenerating livers lacking SOCS1 in hepatocytes but not in SOCS3-deficient livers. These results show that some of the negative correlations between SOCS1 or SOCS3 and the oncogenic signaling pathway genes, notably *CDK2* and *AURKA*, observed in the TCGA dataset are recapitulated in SOCS1- or SOCS3-deficient murine hepatocytes during physiological and pathological hepatocyte proliferation. Consistent with these results, TCGA-LIHC specimens displaying low SOCS1/high CDK2 and low SOCS3/high AURKA expression displayed poor prognosis compared to low SOCS1/low CDK2 and low SOCS3/low AURKA groups (Figure 8D).

### 3.8. Impact of oncogenic signaling genes on the predictive potential of SOCS1 and SOCS3

Next, we carried out a systematic analysis of the predictive potential of *SOCS1* and *SOCS3* when combined with the expression levels of each of the oncogenic signaling pathway genes using the Cox proportional hazards model (Table 2). Even though a high expression of many oncogenic signaling pathway genes independently predicted poor survival (Supplementary Table 1) and some of them showed a better prognostic potential when combined with SOCS1/SOCS3 (*CDK2* and *AURKA*; Figure 8D), the Cox model revealed a different set of genes with prognostic potential (Table 2). Notably, low *SOCS1* displayed a significant predictive potential when combined with *CDK1, CXCL8, CSF1, DAB2* and *TSC1*. Among them, *CXCL8* and *DAB2* also predicted poor survival in combination with low *SOCS3*, even though the latter did not display independent predictive ability. On the other hand, high *SOCS1* expression showed limited synergy with most other genes in predicting better survival (Supplementary Table 1, Table 2). Notably, high *SOCS1* even lost its good prognosis in tumors with high *E2F7*, which independently predicts poor survival, or with low *OPCML*, which is a tumor suppressor [52] but has no predictive ability of its own. A similar scenario was observed for *SOCS3* (Table 2). Moreover, only very few genes that show inverse expression levels with SOCS1/SOCS3 (*PIK3R1, AKT1S1, AUKRA*) are represented in the Cox model (Tables 1 and 2). These observations reveal the complexities of using *SOCS1/SOCS3* as predictive biomarkers as they seem to impact on and be influenced by diverse oncogenic pathway genes and highlight the need for testing such predictions in different clinical study cohorts in order to develop a consensus biomarker panel for prognostication and to identify new drug targets.

**Table 2.**
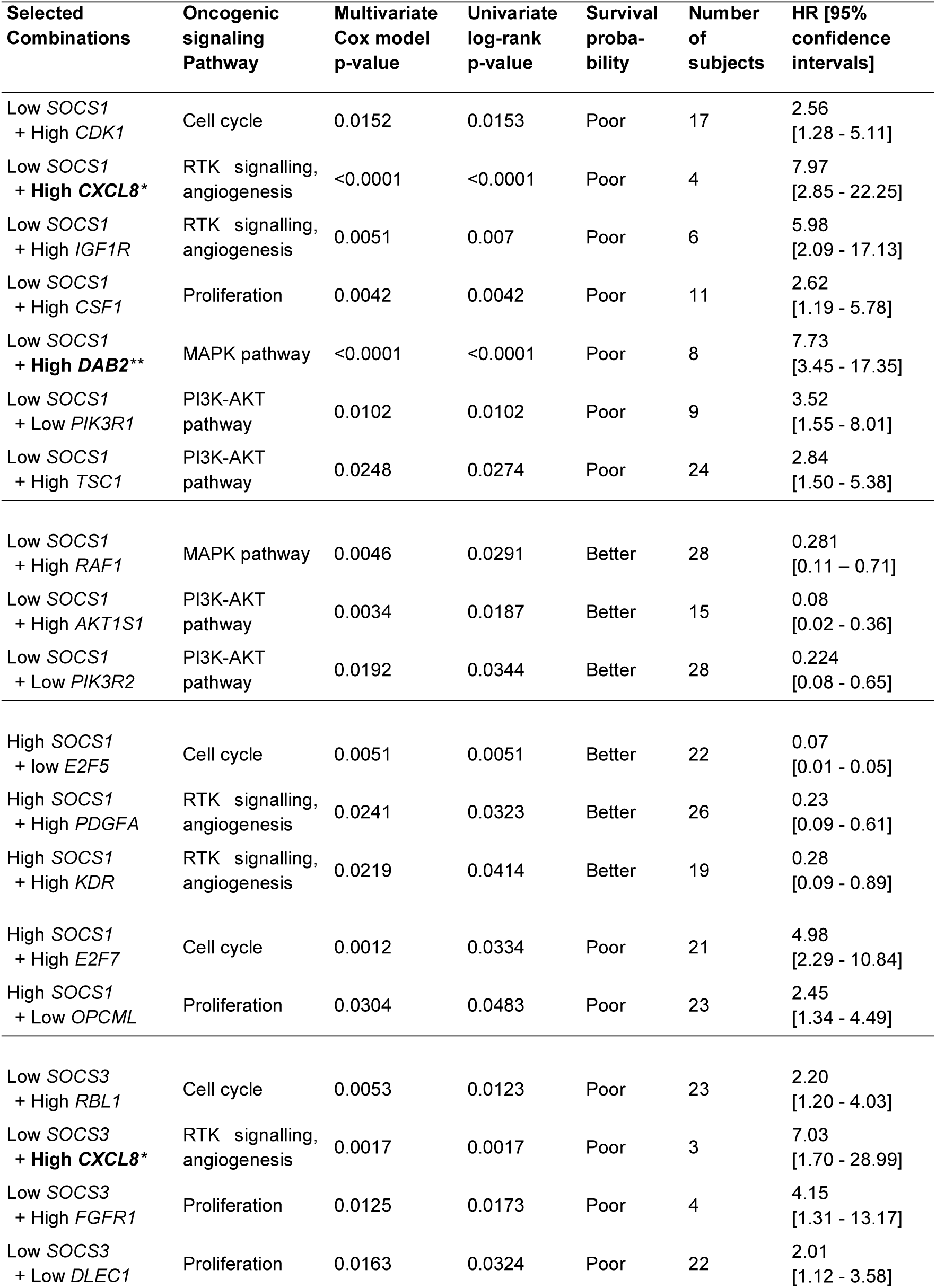
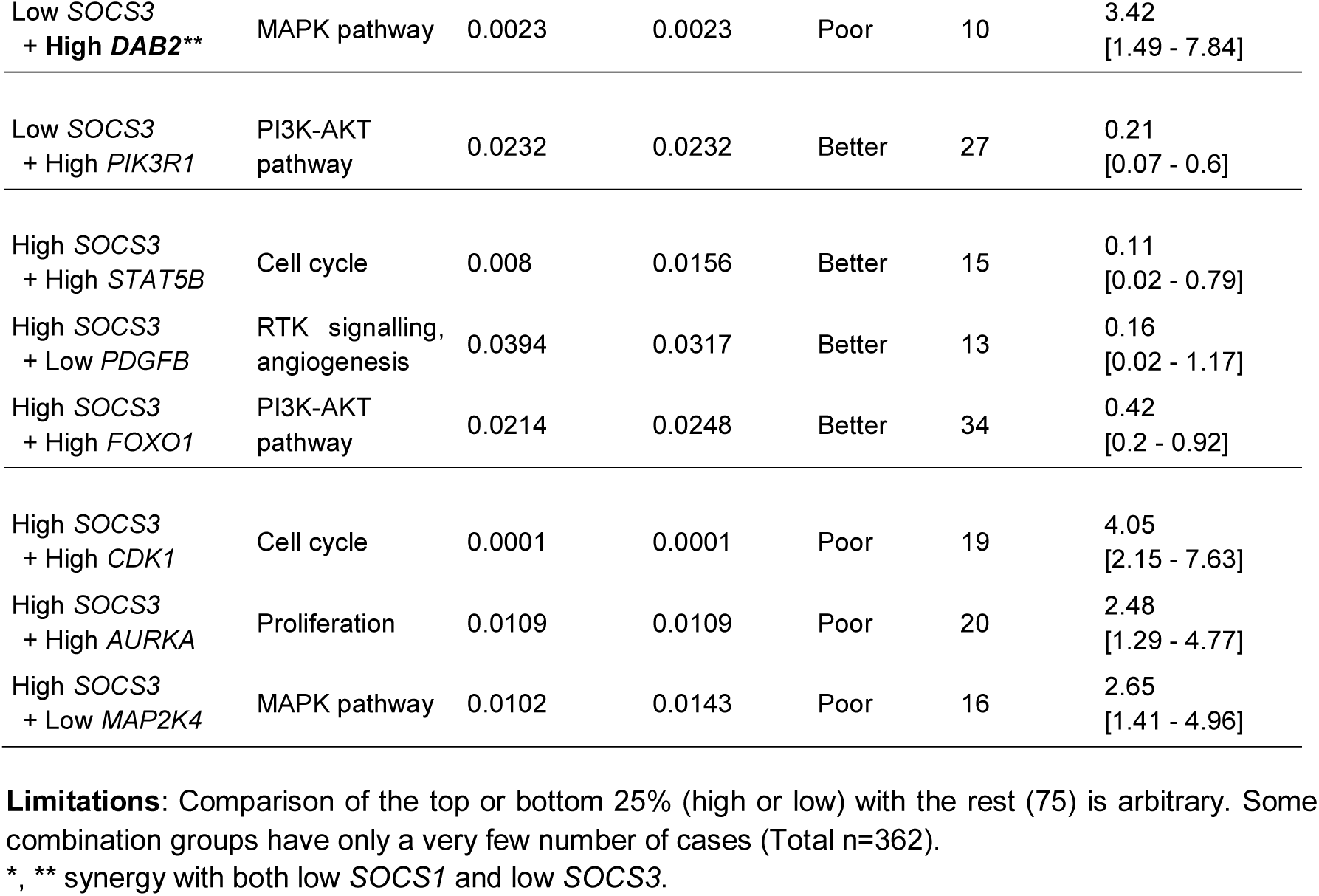
Combinations of *SOCS1* or *SOCS3* and oncogenic signalling pathway genes that show significant prognosis in the Cox proportional hazard model.

| Selected Combinations | Oncogenic signaling Pathway | Multivariate Cox model p-value | Univariate log-rank p-value | Survival probability | Number of subjects | HR [95% confidence intervals] |
| --- | --- | --- | --- | --- | --- | --- |
| Low *SOCS1* + High *CDK1* | Cell cycle | 0.0152 | 0.0153 | Poor | 17 | 2.56<br>[1.28 - 5.11] |
| Low *SOCS1* + High *CXCL8* * | RTK signalling, angiogenesis | <0.0001 | <0.0001 | Poor | 4 | 7.97<br>[2.85 - 22.25] |
| Low *SOCS1* + High *IGF1R* | RTK signalling, angiogenesis | 0.0051 | 0.007 | Poor | 6 | 5.98<br>[2.09 - 17.13] |
| Low *SOCS1* + High *CSF1* | Proliferation | 0.0042 | 0.0042 | Poor | 11 | 2.62<br>[1.19 - 5.78] |
| Low *SOCS1* + High *DAB2* ** | MAPK pathway | <0.0001 | <0.0001 | Poor | 8 | 7.73<br>[3.45 - 17.35] |
| Low *SOCS1* + Low *PIK3R1* | PI3K-AKT pathway | 0.0102 | 0.0102 | Poor | 9 | 3.52<br>[1.55 - 8.01] |
| Low *SOCS1* + High *TSC1* | PI3K-AKT pathway | 0.0248 | 0.0274 | Poor | 24 | 2.84<br>[1.50 - 5.38] |
| Low *SOCS1* + High *RAF1* | MAPK pathway | 0.0046 | 0.0291 | Better | 28 | 0.281<br>[0.11 - 0.71] |
| Low *SOCS1* + High *AKT1S1* | PI3K-AKT pathway | 0.0034 | 0.0187 | Better | 15 | 0.08<br>[0.02 - 0.36] |
| Low *SOCS1* + Low *PIK3R2* | PI3K-AKT pathway | 0.0192 | 0.0344 | Better | 28 | 0.224<br>[0.08 - 0.65] |
| High *SOCS1* + low *E2F5* | Cell cycle | 0.0051 | 0.0051 | Better | 22 | 0.07<br>[0.01 - 0.05] |
| High *SOCS1* + High *PDGFA* | RTK signalling, angiogenesis | 0.0241 | 0.0323 | Better | 26 | 0.23<br>[0.09 - 0.61] |
| High *SOCS1* + High *KDR* | RTK signalling, angiogenesis | 0.0219 | 0.0414 | Better | 19 | 0.28<br>[0.09 - 0.89] |
| High *SOCS1* + High *E2F7* | Cell cycle | 0.0012 | 0.0334 | Poor | 21 | 4.98<br>[2.29 - 10.84] |
| High *SOCS1* + Low *OPCML* | Proliferation | 0.0304 | 0.0483 | Poor | 23 | 2.45<br>[1.34 - 4.49] |
| Low *SOCS3* + High *RBL1* | Cell cycle | 0.0053 | 0.0123 | Poor | 23 | 2.20<br>[1.20 - 4.03] |
| Low *SOCS3* + High *CXCL8* * | RTK signalling, angiogenesis | 0.0017 | 0.0017 | Poor | 3 | 7.03<br>[1.70 - 28.99] |
| Low *SOCS3* + High *FGFR1* | Proliferation | 0.0125 | 0.0173 | Poor | 4 | 4.15<br>[1.31 - 13.17] |
| Low *SOCS3* + Low *DLEC1* | Proliferation | 0.0163 | 0.0324 | Poor | 22 | 2.01<br>[1.12 - 3.58] |
| Low *SOCS3*<br>+ High *DAB2* ** | MAPK pathway | 0.0023 | 0.0023 | Poor | 10 | 3.42<br>[1.49 - 7.84] |
| Low *SOCS3*<br>+ High *PIK3R1* | PI3K-AKT<br>pathway | 0.0232 | 0.0232 | Better | 27 | 0.21<br>[0.07 - 0.6] |
| High *SOCS3*<br>+ High *STAT5B* | Cell cycle | 0.008 | 0.0156 | Better | 15 | 0.11<br>[0.02 - 0.79] |
| High *SOCS3*<br>+ Low *PDGFB* | RTK signalling,<br>angiogenesis | 0.0394 | 0.0317 | Better | 13 | 0.16<br>[0.02 - 1.17] |
| High *SOCS3*<br>+ High *FOXO1* | PI3K-AKT<br>pathway | 0.0214 | 0.0248 | Better | 34 | 0.42<br>[0.2 - 0.92] |
| High *SOCS3*<br>+ High *CDK1* | Cell cycle | 0.0001 | 0.0001 | Poor | 19 | 4.05<br>[2.15 - 7.63] |
| High *SOCS3*<br>+ High *AURKA* | Proliferation | 0.0109 | 0.0109 | Poor | 20 | 2.48<br>[1.29 - 4.77] |
| High *SOCS3*<br>+ Low *MAP2K4* | MAPK pathway | 0.0102 | 0.0143 | Poor | 16 | 2.65<br>[1.41 - 4.96] |

**Limitations:** Comparison of the top or bottom 25% (high or low) with the rest (75) is arbitrary. Some combination groups have only a very few number of cases (Total n=362).
\*, \*\* synergy with both low *SOCS1* and low *SOCS3*.

## 4. Discussion

Our study has revealed notable differences between published reports on the prognostic utility of specific oncogenic signaling pathway genes in HCC from monocentric studies on the one hand, and the multicentric TCGA-LIHC dataset on the other. For example, epigenetic repression of the *SOCS1* gene reported in several studies in up to 65% of HCC specimens is not reflected in *SOCS1* mRNA expression within the TCGA dataset, whereas *SOCS3* gene reported being repressed only in 33% of HCC cases showed downregulation in the TCGA data [9-11]. The reasons for this apparent discrepancy between TCGA-LIHC dataset and other published reports on *SOCS* gene expression in HCC are unclear. Whereas methylation data based on a positive PCR product reflects the repressed status of the gene in hepatocytes, it is possible that induction of the *SOCS* gene expression in liver-resident and infiltrating immune cells and hepatic stromal cells by myriad of cytokines and growth factors might contribute to the overall *SOCS1* transcript levels in the TCGA dataset. In support of this possibility, *SOCS1* mRNA expression positively correlated with *CD247* (CD3 zeta chain, all T cells), *CD8A* (CD8^+^ T cells), *NCAM1* (CD56, NK cells) and *IFNG* (activated T and NK cells) (data not shown). Nonetheless, even though *SOCS1* mRNA expression was not significantly different between tumor and normal tissues, higher transcript levels strongly correlated with patient survival (Figure 1C), highlighting the potential prognostic utility of *SOCS1* expression in HCC.

The positive correlations in the expression of *SOCS1*/*SOCS3* and the oncogenic signaling pathway genes may result from the induction of intact *SOCS1* and *SOCS3* genes as a negative feedback loop to control oncogenic signaling. On the other hand, the negative correlations could arise either from increased oncogenic signaling due to reduced *SOCS1/SOCS3* expression, or from reduced oncogenic signaling due to increased *SOCS1/SOCS3* expression. Moreover, SOCS1 and SOCS3 might target certain transcriptional activators or repressors as substrate specific adaptors in ubiquitin-mediated proteasomal degradation. Thus, the loss of SOCS1 or SOCS3 expression may also affect gene expression indirectly. In the following paragraphs, we discuss the impact of genes that show a negative correlation with *SOCS1* and *SOCS3* on HCC pathogenesis.

### 4.1. Cell Cycle Regulation

As in other cancers, uncontrolled cell cycle progression is a key feature of HCC that results from aberrant expression of cell cycle proteins and/or their regulators [53]. Among the cell cycle genes that showed a negative correlation with *SOCS1/SOCS3, CDK2* and *E2F8* are independent predictors of poor survival (Figure 2B). After the initiation of the cell cycle by growth factor-induced activation of cyclin D1/CDK4/6 complexes, the cyclin E/CDK2 activity induced at the G1/S phase boundary is critical for phosphorylating the retinoblastoma protein (RB) and to relieve E2F transcription factors from inhibition to induce the other genes needed for cell cycle progression [53]. Under conditions of increased cyclin D1 availability, for example from increased growth factor signaling, CDK2 promotes hepatocyte proliferation. CDK2 is needed for the initiation of HCC, although not for progression as HCC may acquire CDK2 independence [54-56]. CDK2 and other CDKs are considered potential therapeutic targets in many cancers including HCC [57]. While high *CDK2* expression correlated with poor survival in our study, Sonntag et al., [55] did not find significant prognostic value for *CDK2*, possibly because the latter used the median value to separate the low and high expression groups, whereas in our study we compared the high one-quartile group and the remaining with low/medium expression.

The E2F family transcription factors are critical regulators of the cell cycle and include transcriptional activators (E2F1-3) and transcriptional repressors, which are either typical (E2F4-6) or atypical (E2F7,8). However, the atypical E2F8 is also implicated in promoting E2F1 transcription in HCC pathogenesis and is considered a potential therapeutic target [58-60]. High *E2F8* transcript levels inversely correlate with *SOCS3* expression and predict poor survival (Figure 2C). It is noteworthy that the expression of *CDK2* and *E2F8* are significantly increased during HCC progression (Supplementary Figure S2B). Understanding how SOCS1 and SOCS3 influence *CDK2* and *E2F8* expression, respectively, could lead to finding ways to block their oncogenic activities in HCC.

Other cell cycle genes *STAT5B, CDK6, RBL2, CCND1, CDKN1B* and *E2F1* that are inversely correlated to *SOCS1/SOCS3* expression are implicated in HCC initiation, progression or both in preclinical models that are also supported by clinical data [61-66]. However, none of these genes show independent prognostic value within the TCGA-LIHC dataset. On the other hand, several cell cycle pathway genes with either positive correlation or no correlation to *SOCS1* displayed independent ability to predict poor survival (Supplementary Table 1, Figure 7). Multivariate analysis of all the cell cycle genes showed that high *CDK1* expression displayed synergistic predictive potential with low *SOCS1*, or even with high *SOCS3* levels (Table 2). It is noteworthy that CDK1 promotes activation of the β-catenin signaling pathway in cancer stem cells and that inhibiting CDK1 increased the efficacy of Sorafenib in a preclinical model [67]. Whether low *SOCS1* and high *CDK1* together predict aggressive HCC that would be responsive to such targeted therapy could be assessed, even retrospectively, in human study cohorts.

### 4.2. RTK signaling and angiogenesis pathways

RTK signaling in hepatocytes and endothelial cells is a key promoter of HCC pathogenesis [2,33,68]. Indeed, deregulation of this pathway by genomic alterations is over-represented in the TCGA-LIHC dataset, and drugs targeting this pathway such as Sorafenib and Regorafenib are already being used or in advanced clinical trials [2,6,69]. However, it is widely perceived that drug choice based on biomarker analysis could improve the treatment outcome. Key RTK signaling/angiogenesis genes *MET, ERBB2, KDR* and *EGFR*, all implicated in HCC [36,70-73], negatively correlate with *SOCS1*, and the first two with *SOCS3* as well. Deregulated EGFR and KDR signaling can contribute to Sorafenib resistance in advanced HCC [72,74]. However, none of these four genes were able to independently predict patient survival within the TCGA-LIHC dataset (Figure 3C). This notion is supported by the failure of MET-targeting therapeutics to improve survival outcome that has been recently attributed, at least partly, to the ability of kinase-inhibited MET to promote cell survival via promoting autophagy [75]. Even though *MET* expression alone was not predictive, ERBB3, which contributes to the resistance to MET inhibition [76] displayed a high predictive potential (Supplementary Table 1). Similarly, even though KDR was not predictive, its ligand *VEGFA* showed a very strong predictive potential (Figure 3C). These findings identify *ERBB3* and *VEGFA* as potential biomarkers for targeted therapies.

CXCL8 (IL-8), a chemokine secreted by inflammatory cells including activated HSCs, induces angiogenic growth factors such as VEGFA in HCC cells and promotes angiogenesis [77-79]. Strikingly, *CXCL8*, which shows a strong negative prognosis in HCC, displayed marked synergy with low *SOCS1* or *SOCS3* in multivariate analysis (Table 2), suggesting the potential use of these markers together. Indeed, CXCL8 receptor (CXCR1, CXCR2) antagonists [80] could be an important addition to targeted therapeutics in HCC.

### 4.3. Other growth factors/proliferation signaling pathways and telomerase maintenance

Among the other growth stimulatory pathways, CSF1-CSFR1 signaling is implicated in promoting HCC via tumor-associated macrophages, and inhibition of this pathway is considered an important strategy for immunotherapy of HCC [39,81]. Indeed, among the other growth factors, only *CSF1* (though not *CSF1R*) displayed a very high predictive value (Supplementary Table 1). Both *SOCS1* and *SOCS3* expression show a high positive correlation with *CSF1* and *CSF1R*. SOCS1 has been shown to interact with CSF1R and regulate its signaling pathways in cellular systems [82] that are captured in the STRING pathway, which also indicates that SOCS1 and SOCS3 could also be indirectly linked to CSF1 (Supplementary Figure 4A). The IGF1-IFG1R system is implicated in the pathogenesis of many cancers. In HCC, IGF1 is downregulated and IGF2 and IGF1R are upregulated, and the latter can form heterodimeric receptors with insulin receptor subunits that bind IGF2 to provide mitogenic signals [42]. However, *IGF1R* did not have a prognostic capacity. Similarly, FGFR1 signaling, which contributes to resistance to MET inhibition [41], also had no independent predictive value. The only gene within the other proliferation signaling pathway that showed prognostic potential was *AURKA*, which promotes centrosome duplication and cytokinesis at the G2/M phase of the cell cycle. *AURKA* is a biomarker for HCC development and progression and is a potential target for therapy [83,84]. *AURKA* expression is dramatically high in TCGA-LIHC dataset, with a significantly increased expression as the disease progresses, and displays a strong predictive ability for disease outcome (Figure 4B, 4C, Supplementary Figure S4B, Tables 1 and 2). *AURKA* expression is negatively correlated to *SOCS3*, even though STRING analysis does not reveal any relationship between the two proteins (Supplementary Fig S4A). The negative regulation of AURKA by SOCS3 is also highlighted by a more than 100-fold increase in the expression of *Aurka* in the regenerating livers of hepatocyte-specific SOCS3-deficient mice, and significant upregulation of this gene in liver tumors induced in these mice (Figure 8A, 8B). One possible mechanism by which SOCS3 could modulate AURKA expression could be via p53, which represses *AURKA* [85]. Indeed, an earlier study reported frequent AURKA overexpression in HCC that was associated with p53 mutation [86]. SOCS3 can promote transcriptional activation of p53 [22]. Whether SOCS3 can also modulate the repressive function of p53 is not known. It is noteworthy that high *AURKA* expression can occur even with high *SOCS3* expression, which together predict poor prognosis in multivariate analysis (Table 2). However, SOCS1, which was shown to activate p53 earlier [87,88], did not correlate with *AURKA* expression in the TCGA dataset. Clearly, further studies are needed to elucidate the mutual exclusivity of *SOCS3* and *AURKA* expression in HCC.

### 4.4. RAS-RAF-MAPK signaling pathways

The RAS-RAF-MAPK pathway is frequently perturbed in HCC and thus is an important therapeutic target [89]. Indeed, the RAF kinase is a key target of Sorafenib that is already used in HCC therapy. Whereas BRAF mutations are commonly found in cholangiocarcinoma but not in HCC, immunohistochemical analysis of HCC specimens in a Chinese cohort identified RAF1 but not ERK to have a prognostic value [90,91]. On the other hand, our study on the TCGA-LIHC dataset revealed that *BRAF* as well as *MAPK1* (ERK2), which show mutual exclusivity with both *SOCS1* and *SOCS3*, display significant predictive value (Figure 5). Similarly, *MAP3K4* (MEKK4), which is inversely correlated to SOCS3 expression, is also highly predictive of poor survival.

Several genes of the RAS-RAF-MAPK pathway that are coordinately regulated with *SOCS1* and/or *SOCS3* also displayed strong predictive ability in the TCGA dataset (Supplementary Table 1). Of these, those involved in the negative regulation of the RAS-RAF-MAPK pathway have been reported to have a predictive potential [47,48,92]. Members of the RAS association domain family (RASSF) of RAS inhibitors (RASSF1A, RASSF2A and RASSF5 (NORE1A, NORE1B) that inhibit RAS activity), RASAL1 that inhibits RAS activity by promoting the intrinsic GTPase activity, and SPRED1 and SPRED2 that inhibit RAF kinases are frequently repressed by promoter methylation in HCC [47,48,92]. DAB2, which attenuates the RAS activation downstream of RTK signaling by binding to GRB2 and dissociating its interaction with SOS1, is also repressed by promoter methylation [47] and showed coordinate regulation with *SOCS1* and *SOCS3*. As promoter hypermethylation also represses *SOCS1* and *SOCS3*, it is possible that epigenetic repression of both *SOCS* genes as well as the endogenous negative regulators the RAS-RAF-MAPK pathway likely contributes to their coordinate regulation that amplifies the proliferation and anti-apoptotic functions of this way, contributing to HCC pathogenesis.

Surprisingly, high expression of *RASSF1* and *DAB2* predicted poor survival in the TCGA-HCC dataset (Supplementary Table 1), instead of a better prognosis expected of their function as negative regulators of the RAS-MAPK pathway. High *DAB2* expression within low *SOCS1* or *SOCS3* expressing subgroups also predicted poor overall survival (Table 2). The reason for this apparent discrepancy is unclear. It is possible that the upregulation of *RASSF1* and *DAB2* may result from mutations that disrupt the normal functions of these tumor suppressors, as in the case of mutant p53, which is highly expressed in many cancers [93]. However, only a negligible proportion of cases in the TCGA dataset revealed mutations for *RASSF1* or *DAB2* (data not shown). It is equally possible that their increased expression could result from a compensatory increase in response to the increased activity of this pathway or mutations in their target proteins. Clearly further studies are needed to resolve this conundrum.

### 4.5. PI3K-AKT-MTOR Signaling pathway

The MTOR pathway is frequently activated in HCC and is associated with poor prognosis [51]. Our findings reveal that out of eleven genes of the PI3K-AKT-MTOR pathway driver genes analyzed for survival analysis, *RPTOR, PIK3CA, TSC1, MLST8* and *AKT1S1* showed a pronounced negative impact on patient survival whereas *PIK3R1* showed favorable impact (Figure 6C), raising the possibility of using these genes as prognostic markers. Whereas low *PIK3R1* synergized with low *SOCS1* to predict poor survival as expected, paradoxically, cases with low *SOCS1* and high *AKT1S1* (a key substrate of AKT in the MTOR signaling pathway) showed better survival (Table 2) for obscure reasons. Both SOCS1 and SOCS3 have been implicated in regulating the PI3K-AKT pathway upstream of MTOR. By virtue of their ability to promote ubiquitination and proteasomal degradation of insulin receptor substrates 1 and 2 (IRS1, IRS2), which link RTK signaling to PI3K, SOCS1 and SOCS3 can regulate AKT activation in the context of insulin resistance in the liver and other organs [94-96]. We have shown that SOCS1-deficient primary hepatocytes show increased AKT activation in response to HGF [28]. *SOCS3* knockdown in endothelial cells increases proliferation and sprouting through activation of the MTOR pathway [97]. Even though SOCS1/SOCS3-mediated attenuation of AKT activation could result from the inhibition of RTK signaling, the STRING analysis connects SOCS1 to the PI3K-AKT-MTOR pathway via PIK3R1 (Supplementary Figure 6A) that was based on a large-scale yeast two-hybrid screen for phosphor-tyrosine-dependent protein interactions and mass spectrometry [98,99]. Recently, the Fraternali lab has placed SOCS1 and PIK3R1 within a network of many RTKs including MET and ERBB2 through an analysis method called short loop motif profiling [100]. Our findings show that the expression of both *SOCS1* and *SOCS3* show a high degree of mutual exclusivity with *PIK3R1* (Figure 6A). Even though this gene codes for the p85 regulatory subunit of PI3K, there is strong evidence indicating its role also as a tumor suppressor by positively modulating PTEN, a negative regulator of the PI3K pathway, and by modulating AKT and STAT3 activation [101-103]. Consistent with this, high *PIK3R1* predicts favorable survival, whereas its lower expression in synergy with low *SOCS1* predicts poor survival (Table 2). Given the tumor suppressor functions and overlapping mechanisms of action of SOCS1, SOCS3 and PIK3R1, further work is needed to disentangle the highly significant negative correlation between *SOCS1/SOCS3* and *PIK3R1*.

## 5. Conclusions

The most significant outcomes of our study are (i) the prognostic significance of *SOCS1* gene expression in HCC either alone or in combination with certain other genes, and (ii) the discordant data on the predictive potential of certain biomarkers reported earlier within the TCGA-HCC dataset. Clearly, further studies in other cohorts are needed to confirm or contradict these findings. Besides, we observed coordinated expression of several oncogenic signaling pathway genes and *SOCS1/SOCS3*, presumably reflecting activation of negative feedback loops. However, nearly half of the PI3K-AKT-MTOR pathway genes showed mutual exclusivity with *SOCS1/SOCS3*, suggesting the loss of SOCS-dependent regulation of RTKs contributing to the increased activity of this signaling pathway. Finally, our study identified at least three genes, *RASSF1* and *DAB2* in the RAS-RAF-MAPK pathway and *PIK3R1* in the PI3K-AKT-MTOR pathway that showed a predictive value opposite of their expected functions, indicating a need for further investigations. Collectively, *SOCS1* and certain key genes of the oncogenic signaling pathways that show high predictive value in this study could be developed as combination biomarkers for patient-oriented precision therapeutics in HCC.

## Supporting information

MGM-Khan_SOCS1,3_TCGA-LIHC_V1-Supplemnet

## Author Contributions

Conceptualization, M.K., S.R., and S.I.; Data Analysis and interpretation, M.K., M.R., B.V., A.I., and A.G.; Experiments, M.K., and A.G.; Manuscript drafting and editing, M.K., M.R., S.R., and S.I. All authors have read the manuscript and approved the final version.

## Acknowledgments

MK was supported by doctoral scholarship from FRQNT. MR is supported by the ‘Abdenour-Nabid’ and AI by the ‘VoiceAge’ graduate fellowships of the Faculty of Medicine, Université de Sherbrooke. AG is a recipient of postdoctoral fellowship from FRQS. We thank Ms Marie-Pierre Garant of the CRCHUS for the Cox proportional hazard analysis.

## Supplementary Material

**Figure S1.** Workflow of this study.

**Figure S2.** Cell cycle regulation genes: interactions with SOCS1 and SOCS3, and expression across the tumor grades in TCGA-LIHC dataset.

**Figure S3.** RTK signaling and angiogenesis pathway genes: interactions with SOCS1 and SOCS3, and expression across the tumor grades in TCGA-LIHC dataset.

**Figure S4.** Other growth signaling pathway genes: interactions with SOCS1 and SOCS3, and expression across the tumor grades in TCGA-LIHC dataset.

**Figure S5.** RAS-RAF-MEK-MAPK pathway genes: interactions with SOCS1 and SOCS3, and expression across the tumor grades in TCGA-LIHC dataset.

**Figure S6.** PI3K-AKT-MTOR pathway genes: interactions with SOCS1 and SOCS3, and expression across the tumor grades in TCGA-LIHC dataset.

**Table S1.** Impact of the expression of oncogenic pathway genes on survival probability in the TCGA-LIHC dataset.

