## Supplementary material for "Prognostic significance of *SOCS1* and *SOCS3* tumor suppressors in hepatocellular carcinoma and its correlation to key oncogenic signaling pathways": MGM-Khan_SOCS1,3_TCGA-LIHC_V1-Supplemnet

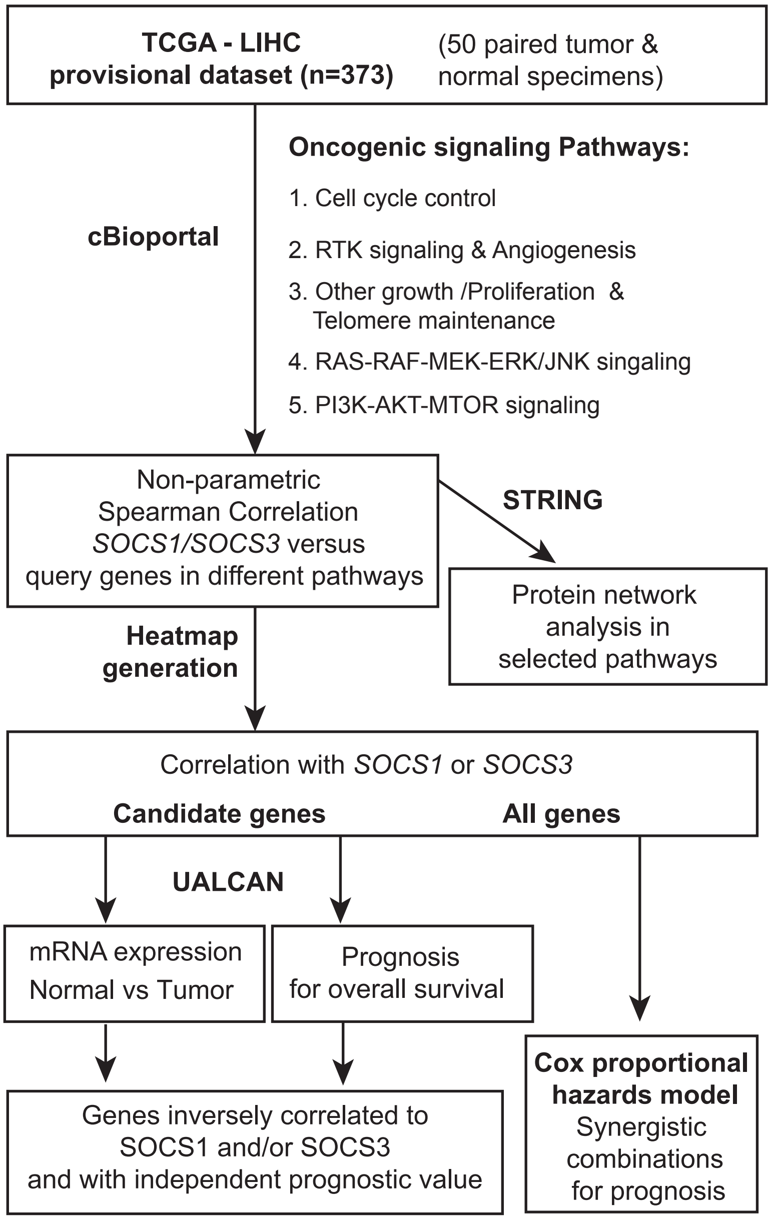


**Figure S1. Workflow of this study.**

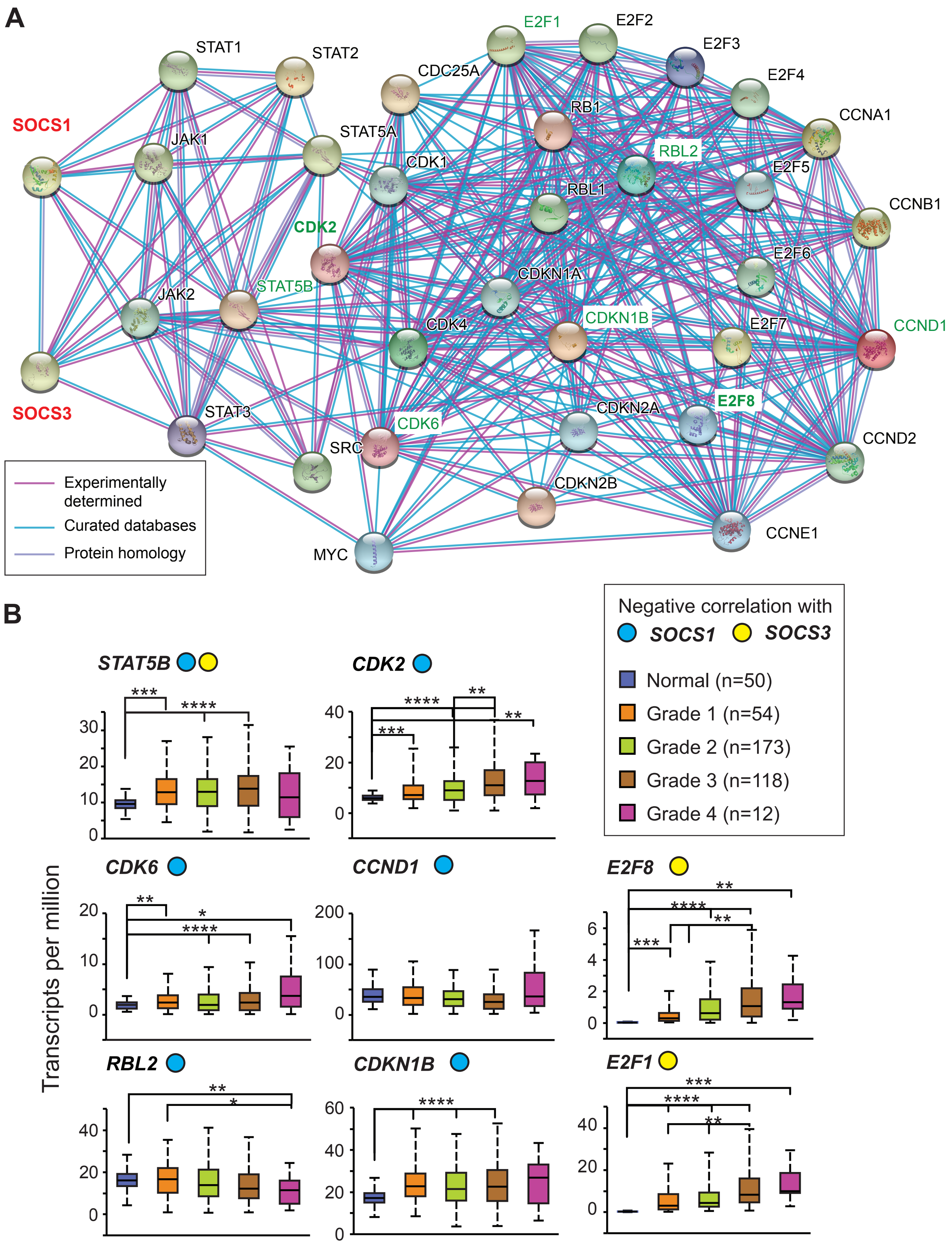


**Figure S2. Cell cycle regulation genes: interactions with SOCS1 and SOCS3, and expression across the tumor grades in TCGA-LIHC dataset.** (A) Protein-protein interaction network between SOCS1, SOCS3 and proteins of the cell cycle regulation pathway implicated in oncogenesis evaluated by the STRING analysis tool. Only medium and high confidence interactions with score above 0.40, supported by experiments and/or curated databases on signal transduction pathways were included. (B) Genes that show significant negative correlation with SOCS1 and/or SOCS3 (Figure 3A, 3B) were evaluated for their expression levels in different grades of HCC tumors compared to normal liver tissues.

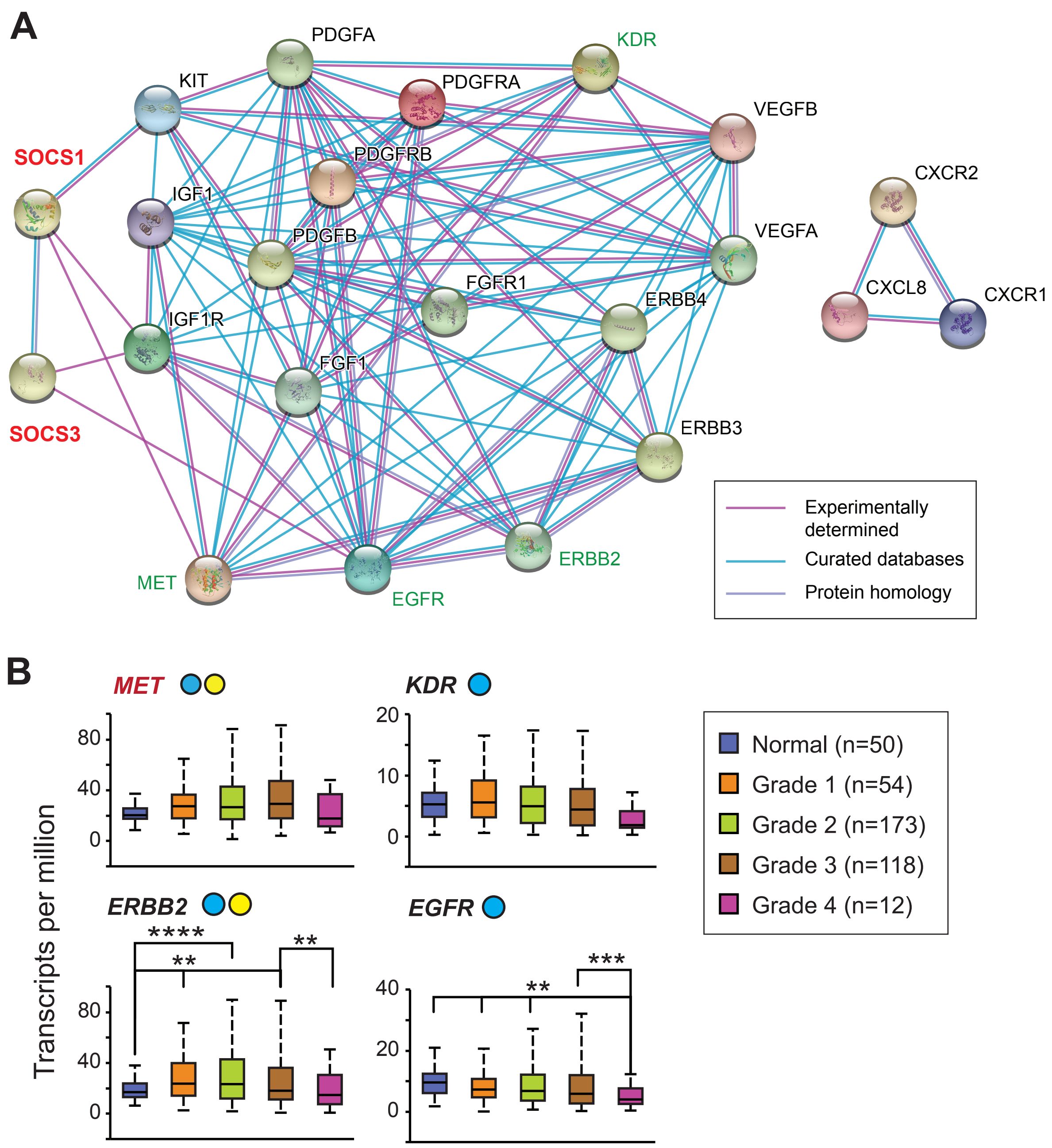


**Figure S3.** RTK signaling and angiogenesis pathway genes: interactions with SOCS1 and SOCS3, and expression across the tumor grades in TCGA-LIHC dataset.

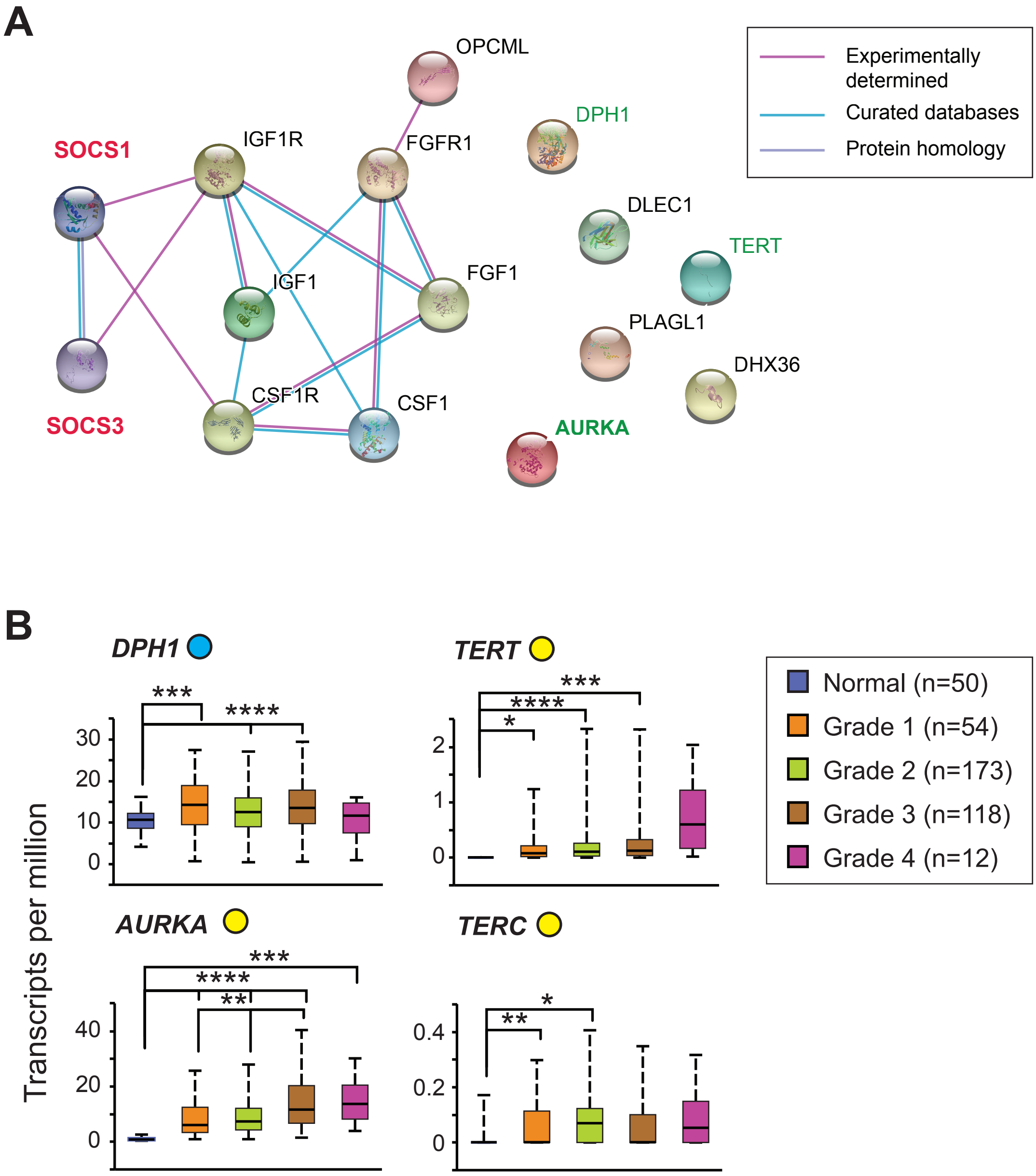


**Figure S4.** Other growth signaling pathway genes: interactions with SOCS1 and SOCS3, and expression across the tumor grades in TCGA-LIHC dataset.

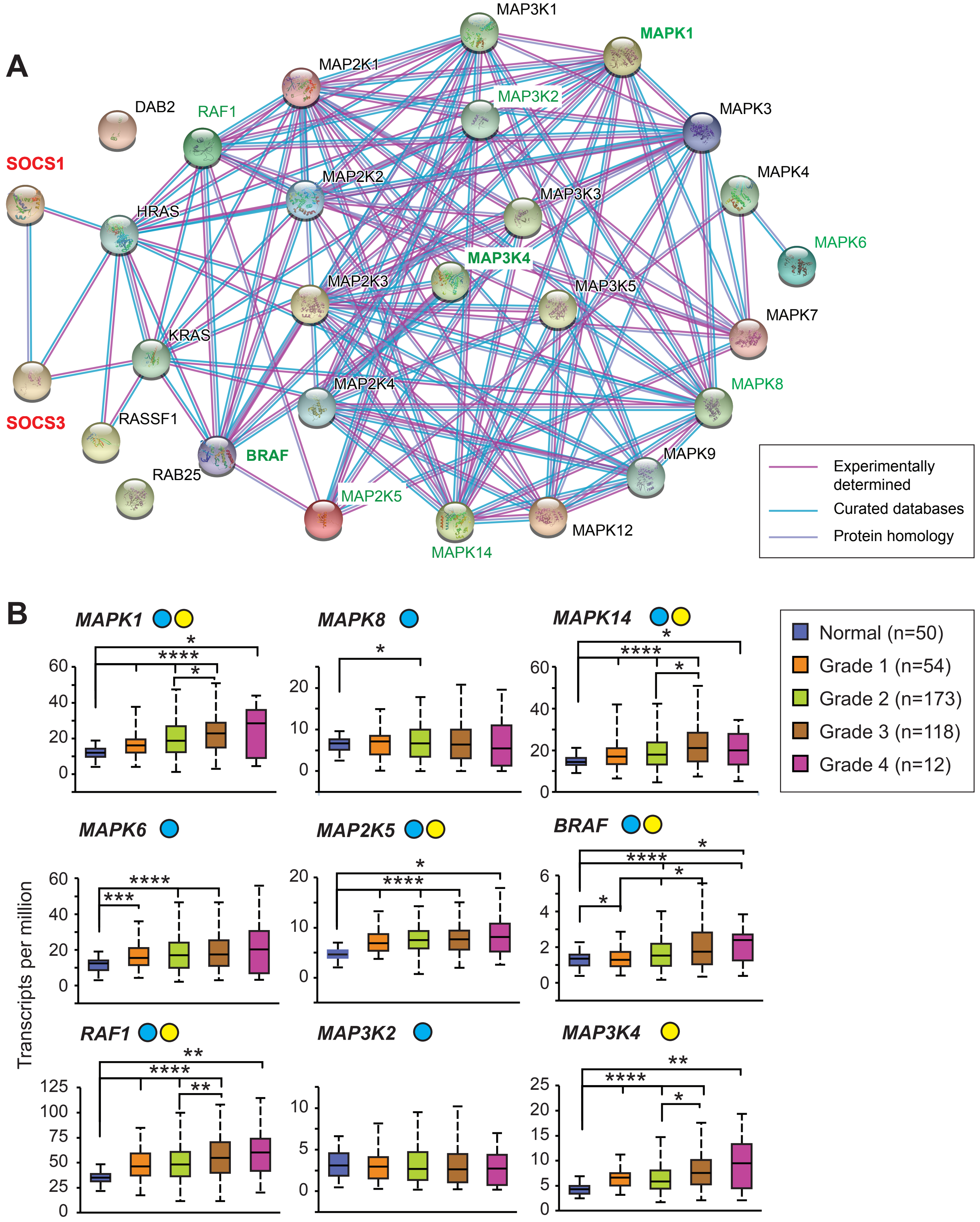


**Figure S5.** RAS-RAF-MEK-MAPK pathway genes: interactions with SOCS1 and SOCS3, and expression across the tumor grades in TCGA-LIHC dataset.

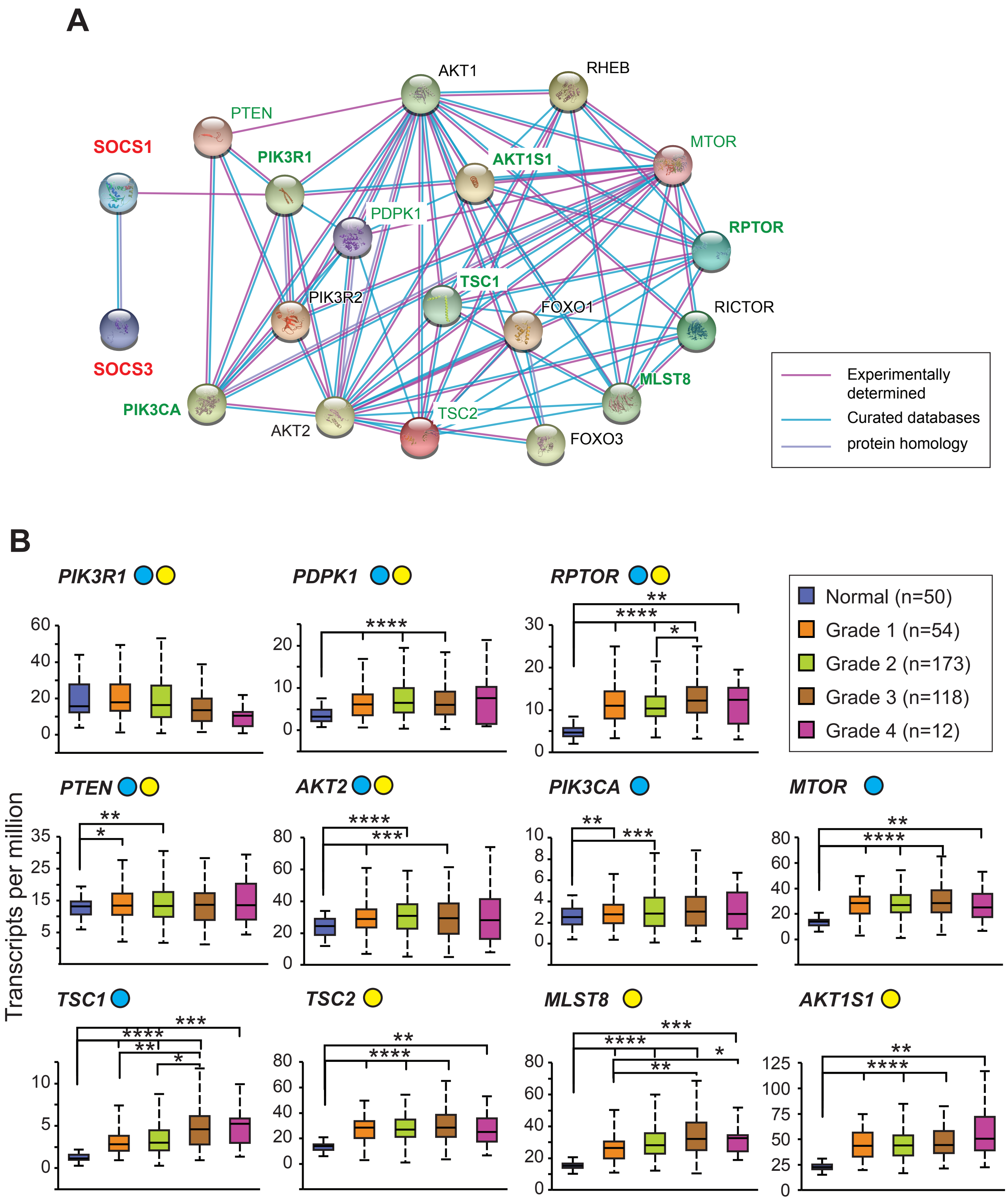


**Figure S6.** PI3K-AKT-MTOR pathway genes: interactions with SOCS1 and SOCS3, and expression across the tumor grades in TCGA-LIHC dataset.

**Supplementary Table 1.** Impact of the expression of oncogenic pathway genes on survival probability in the TCGA-LIHC dataset.

| Oncogenic signaling pathway | Gene Name* | High exp. (n) | Low/Med. Exp (n) | Survival probability p-value | Poor prognosis for expression: |
|---|---|---|---|---|---|
| **Cell Cycle** | *STAT5B* | 92 | 273 | 0.96 | |
| | *CDK6* | 90 | 275 | 0.22 | |
| | *RBL2* | 93 | 272 | 0.76 | |
| | ***CDK2*** | 90 | 275 | **0.011** | High |
| | *CCND1* | 90 | 275 | 0.95 | |
| | *CDKN1B* | 92 | 273 | 0.18 | |
| | *RB1* | 90 | 275 | 0.077 | |
| | ***RBL1*** | 91 | 274 | **0.044** | High |
| | ***E2F7*** | 92 | 273 | **0.0044** | High |
| | ***E2F8*** | 91 | 274 | **0.006** | High |
| | *JAK1* | 91 | 274 | 0.17 | |
| | ***E2F6*** | 89 | 276 | **0.00021** | High |
| | *MYC* | 91 | 274 | 0.32 | |
| | *E2F1* | 92 | 273 | 0.1 | |
| | ***CDK1*** | 89 | 276 | **<0.0001** | High |
| | ***CDC25A*** | 89 | 276 | **<0.0001** | High |
| | ***CDKN2B*** | 91 | 274 | **0.0016** | High |
| | ***E2F5*** | 89 | 276 | **0.0021** | High |
| | ***CCNB1*** | 90 | 275 | **<0.0001** | High |
| | ***CDK4*** | 88 | 277 | **<0.0001** | High |
| | ***SRC*** | 92 | 273 | **0.00049** | High |
| | ***CDKN2A*** | 91 | 274 | **0.014** | High |
| | *STAT2* | 91 | 274 | 0.49 | |
| | *CCNA1* | 92 | 273 | 0.33 | |
| | *E2F3* | 89 | 276 | 0.13 | |
| | *E2F2* | 90 | 275 | 0.096 | |
| | ***E2F4*** | 88 | 277 | **0.00019** | High |
| | ***CCNE1*** | 92 | 273 | **0.00077** | High |
| | *STAT1* | 91 | 274 | 0.16 | |
| | *CDKN1A* | 91 | 274 | 0.75 | |
| | *JAK2* | 90 | 275 | 0.91 | |
| | *STAT5A* | 91 | 274 | 0.75 | |
| | *STAT3* | 92 | 273 | 0.42 | |
| | *CCND2* | 90 | 275 | 0.66 | |
| **RTK signaling & Angiogenesis** | *MET* | 93 | 272 | 0.64 | |
| | *ERBB2* | 93 | 272 | 0.34 | |
| | *KDR* | 93 | 272 | 0.088 | |
| | *EGFR* | 93 | 272 | 0.54 | |
| | ***VEGFA*** | 90 | 275 | **<0.0001** | High |
| | *KIT* | 91 | 274 | 0.093 | |

| | | | | | |
|---|---|---|---|---|---|
| | *PDGFB* | 92 | 273 | 0.96 | |
| | *IGF1* | 93 | 272 | 0.089 | |
| | *PDGFRB* | 92 | 273 | 0.35 | |
| | *ERBB4* | 92 | 273 | 0.67 | |
| | ***ERBB3*** | 92 | 273 | **0.0081** | High |
| | *PDGFA* | 92 | 273 | 0.62 | |
| | *FGF1* | 92 | 273 | 0.46 | |
| | *IGF1R* | 89 | 276 | 0.29 | |
| | *FGFR1* | 92 | 273 | 0.44 | |
| | *PDGFRA* | 91 | 274 | 0.24 | |
| | *VEGFB* | 91 | 274 | 0.61 | |
| | *CXCR1* | 91 | 274 | 0.52 | |
| | *CXCR2* | 91 | 274 | 0.96 | |
| | ***CXCL8*** | 92 | 273 | **0.01** | High |
| **Other growth & Proliferation/TERT** | *DPH1* | 92 | 273 | 0.67 | |
| | *DLEC1* | 92 | 273 | 0.24 | |
| | ***AURKA*** | 90 | 275 | **0.0016** | High |
| | *IGF1* | 93 | 272 | 0.089 | |
| | *OPCML* | 92 | 273 | 0.7 | |
| | *FGF1* | 92 | 273 | 0.46 | |
| | *PLAGL1* | 89 | 276 | 0.74 | |
| | *IGF1R* | 89 | 276 | 0.29 | |
| | *FGFR1* | 92 | 273 | 0.44 | |
| | ***CSF1*** | 91 | 274 | **<0.0001** | High |
| | *CSF1R* | 90 | 275 | 0.15 | |
| | *TERT* | 91 | 274 | 0.51 | |
| | *TERC* | 92 | 273 | 0.31 | |
| **RAS-RAF-MEK-ERK/JNK** | ***MAPK1*** | 91 | 274 | **0.013** | High |
| | *MAPK8* | 93 | 272 | 0.13 | |
| | *MAPK14* | 91 | 274 | 0.19 | |
| | *MAPK6* | 90 | 275 | 0.14 | |
| | *MAP2K5* | 92 | 273 | 0.93 | |
| | ***BRAF*** | 92 | 273 | **0.0066** | High |
| | *RAF1* | 91 | 274 | 0.17 | |
| | *MAP3K2* | 93 | 272 | 0.81 | |
| | ***MAP3K4*** | 89 | 276 | **0.0025** | High |
| | *KRAS* | 93 | 272 | 0.11 | |
| | *MAP2K4* | 92 | 273 | 0.51 | |
| | *MAP3K1* | 90 | 275 | 0.28 | |
| | *MAPK9* | 90 | 275 | 0.21 | |
| | *MAPK4* | 93 | 272 | 0.56 | |
| | *MAP3K3* | 92 | 273 | 0.14 | |
| | *MAP2K1* | 91 | 274 | 0.28 | |
| | *MAP2K3* | 91 | 274 | 0.22 | |
| | ***MAPK7*** | 91 | 274 | **0.0061** | High |
| | ***MAP2K2*** | 90 | 275 | **0.0034** | High |

| | | | | | |
|---|---|---|---|---|---|
| | *RAB25* | 93 | 272 | 0.51 | |
| | ***MAPK12*** | 92 | 273 | **0.026** | High |
| | ***MAPK3*** | 92 | 273 | **0.00015** | High |
| | ***HRAS*** | 88 | 277 | **0.00073** | High |
| | ***RASSF1*** | 89 | 276 | **0.013** | High |
| | *MAP3K5* | 92 | 273 | 0.54 | |
| | ***DAB2*** | 92 | 273 | **0.00046** | High |
| **PI3K-AKT-MTOR** | ***PIK3R1*** | 93 | 272 | **0.012** | Low |
| | *PDPK1* | 93 | 272 | 0.25 | |
| | ***RPTOR*** | 90 | 275 | **0.0043** | High |
| | *PTEN* | 91 | 274 | 0.47 | |
| | *AKT2* | 92 | 273 | 0.38 | |
| | ***PIK3CA*** | 93 | 272 | **0.0088** | High |
| | *MTOR* | 93 | 272 | 0.18 | |
| | ***TSC1*** | 90 | 275 | **0.0028** | High |
| | *TSC2* | 92 | 273 | 0.2 | |
| | ***FOXO1*** | 93 | 272 | **0.027** | Low |
| | ***RICTOR*** | 92 | 273 | **0.0068** | High |
| | ***FOXO3*** | 91 | 274 | **0.0083** | High |
| | ***RHEB*** | 92 | 273 | **0.0026** | High |
| | ***PIK3R2*** | 89 | 276 | **0.018** | High |
| | ***MLST8*** | 89 | 276 | **0.0014** | High |
| | ***AKT1S1*** | 91 | 274 | **0.00073** | High |
| | ***AKT1*** | 90 | 275 | **0.0015** | High |

* Gene names in **boldface**: Significant correlation with *SOCS1* (vary with *SOCS3*; see figures 3-7).

Color codes:

**Green:** negative correlation (mutual exclusivity) with SOCS1 or SOCS3.

**Black:** no correlation with SOCS1.

**Red:** positive correlation (co-occurrence) with SOCS1.

**Orange:** Prognosis discordant from expected functions.
